# Pathogenic Viruses, Genome Integrations, and Viral::Human Chimeric Transcripts Detected by VirusIntegrationFinder Across >30k Human Tumor and Normal Samples

**DOI:** 10.1101/2025.03.10.642430

**Authors:** Brian J. Haas, Anne Van Arsdale, Alexander Dobin, Maxwell Brown, Joshua Gould, Christophe Georgescu, Eleanor Agosta, Sylvain Baulande, Ismail Jamail, Maud Kamal, Ivan Bièche, Jack Lenz, Christina Montagna, Aziz Al’Khafaji

**Affiliations:** Broad Institute of MIT and Harvard, Cambridge MA, 02142, USA; Department of Obstetrics Gynecology and Women’s Health, Albert Einstein College of Medicine, Bronx, NY, 10461, USA; Department of Genetics, Albert Einstein College of Medicine, Bronx, NY, 10461, USA; Cold Spring Harbor Laboratory, New York, NY 11724, USA; Rutgers Cancer Institute of New Jersey, 195 Little Albany St., New Brunswick, NJ, 08901, USA; Institut Curie, PSL University, ICGex Next-Generation Sequencing Platform, 75005 Paris, France; Department of Drug Development and Innovation, Institut Curie, PSL Research University, 75005 Paris & 92210, Saint-Cloud, France; Faculty of Pharmaceutical and Biological Sciences, INSERM U1016, Paris Descartes University, 75005, Paris, France; Rutgers Cancer Institute, 195 Little Albany St., New Brunswick, NJ, 08901, USA

## Abstract

Viruses are a leading cause of human morbidity and mortality. Certain viruses, including human papillomaviruses (HPVs), play a significant role in the etiology of cancer. Detection of viral DNA insertions in the human genome from next generation sequencing data defines viral associations with cancer and other diseases, identifies impacted organs and tissues, provides insights into disease mechanisms and has the potential to enhance clinical evaluations. In this study, we developed VirusIntegrationFinder (CTAT-VIF), a tool for surveying human genome insertions of various human viruses using both DNA and RNA sequencing data. We applied CTAT-VIF to analyze a dataset of over 30,000 tumor and normal samples, as well as more than 1,000 cancer cell lines. This effort resulted in the compilation of a catalog of over 30,0000 virus-human DNA or RNA junctions at more than 20,000 insertion loci and reassessed viral cancer-insertion hotspots across the human genome. Furthermore, we characterized the functional impacts of insertions with respect to human copy number alterations, effects on the expression of flanking human genes, and the identification of potentially oncogenic chimeric human and human/virus fusion transcripts at insertion loci. In addition to confirming known viral associations with specific tumor types, our study revealed both shared and virus-specific insertion hotspots in addition to variable functional impacts based on virus type. Besides some rare events of interest, we also found evidence for sequencing contamination, which underscores the need for vigilance when studying viral content or genome integrations.

## Introduction

Progress in modern sequencing technologies has provided unparalleled opportunities to explore the genetic composition of biological samples at extraordinary resolution and precision. When deployed to clinical specimens, insights extracted from sequencing data can guide mechanistic studies and ultimately influence clinical decisions in personalized medicine. Highly effective and scalable bioinformatics tools are essential for processing the large volume of sequencing data currently available and to integrate the continuous influx of newly sequenced data sets. The detection of viral DNA or RNA in such data sets is a widely used strategy to establish links between specific viruses and numerous diseases (Flippot et al. 2016; Tang and Larsson 2017). Transcriptome sequencing data is especially valuable for comprehensive biological exploration, offering combined functional and genetic insights, whether obtained at bulk or at single cell resolution. Our Cancer Transcriptome Analysis Toolkit (CTAT, (Haas 2019)) can further leverage these data to reconstruct transcript isoforms, capture evidence of sequence variants, fusion transcripts, splicing aberrations, and copy number alterations, and facilitate downstream analysis and interpretation of identified variants and anomalies relevant to cancer.

A small number of viruses, comprising DNA viruses such as Human Papillomaviruses (HPVs), Epstein-Barr Virus (EBV), Human Kaposi’s sarcoma herpesvirus (KSHV; also known as human herpesvirus 8 (HHV8)), Merkel cell polyomavirus (MCV); reverse transcribed viruses like Hepatitis B Virus (HBV) and Human T-Lymphotropic Virus 1 (HTLV-1) alongside the RNA virus Hepatitis C Virus (HCV), are responsible for as much as 10 to 15% of cancers worldwide (Moore and Chang 2010). Among these viruses, HPV is the primary cause of cervical cancer and also significantly contributes to other anogenital cancers as well as head and neck cancers (Stratton and Culkin 2016; Ferreira 2023; Nelson and Mirabello 2023). Notably, HBV is a major contributor to liver cancer (McGlynn et al. 2021), while EBV is associated with gastric cancer (Saito and Kono 2021). While many viruses typically replicate and function as autonomous, independent genomes in human cells, in a number of diseases, viral DNA integrates into the human genome (Chen et al. 2014; Ye et al. 2023). In particular, retrovirus DNA integration into the host cell genome is an essential element of the viral life cycle (Zhang et al. 2018). However, other viral DNAs can also integrate into the host cell genome, even when such an event is not part of the normal virus life cycle. This occurrence is presumably a consequence of aberrant DNA repair or recombination processes associated with genome instability, frequently observed in various human cancers (Chen et al. 2014; Ye et al. 2023). In this context, we introduce our newly developed VirusIntegrationFinder (CTAT-VIF), specifically designed to detect viral DNA and RNA and provide evidence of viral insertions within human genomes.

Viral genomes, including but not limited to HTLV-1 which requires integration into the host genome as part of its life cycle, are detected as joined viral and human genome sequences in various cancerous and precancerous conditions. The viral genomes integrated into the host human genome vary widely, ranging from subgenomic fragments to full-length genomes and tandemly repeated genome multimers (Akagi et al. 2014; McBride and White 2023; Van Arsdale et al. 2024). Integration of viral DNA can lead to a stable association of viral genes with host cells, thereby impacting the function of neighboring genes and potentially altering gene expression. Additionally, such integrations may contribute to genome instability, including copy number amplifications. In some cases, viral insertions are located at known human oncogenes and can contribute to tumorigenesis. Functional retrovirus DNA insertions typically occur at specific positions two base pairs from the ends of the viral long terminal repeats (LTRs), and software to detect these junctions with human DNA is well established (Wildschutte et al. 2016; Berry et al. 2017; Sherman et al. 2017; Holloway et al. 2019; Bale et al. 2021). However, insertions of DNA from other viruses occur at positions throughout much of their viral genomes, which can complicate the detection of bona fide junctions.

Previous bioinformatics tools have been developed to identify viral insertions into the genome, including ViFi (Nguyen et al. 2018), the more recent FastViFi (Javadzadeh et al. 2022), BatVI (Tennakoon and Sung 2017), VirusBreakend (Cameron et al. 2021), VirusFinder2 (VERSE, (Wang et al. 2015)), and nf-VIF (Kamal et al. 2021). These tools vary in alignment methods, data types utilized (RNA-seq, WGS, hybridization capture), and the range of viruses investigated. Evidence for virus insertions primarily relies on chimeric human/virus reads. The search for these reads parallels the approach used for identifying cognate fusion transcripts where chimeric reads derive from different genes of the human genome. Among top-performing fusion transcript tool (Haas et al. 2019; Creason et al. 2021) are STAR-Fusion (Haas et al. 2019), Arriba (Uhrig 2019), and STAR-SEQR (STAR-SEQR 2019) all leveraging the popular STAR aligner (Dobin et al. 2013), known for its capability to detect chimeric reads. We recently adapted STAR for in silico characterization of fusion transcripts through the development of FusionInspector (Haas et al. 2023). Both STAR-Fusion and FusionInspector, discussed in this context, are integral components of CTAT.

In this study, we further adapted the techniques used by STAR-Fusion and FusionInspector to capture evidence of human viruses and viral genome insertions in cancer transcriptomes. This advancement has been implemented as CTAT-VirusIntegrationFinder (CTAT-VIF, **Figure 1a**). Our goals in developing CTAT-VIF were to enable the analysis of a wide diverse range of human viruses in a single software execution, capture evidence of read coverage across viral genomes, and provide evidence for viral genome insertions into the human genome. These insertions correspond to the genome insertion site or to fusion splice products generated between the virus and human genes at insertion loci (collectively referred to as insertions). These functionalities are coupled with easy to navigate interactive visualizations that depict evidence for virus content and any detected virus insertions. We demonstrate that CTAT-VIF achieves accuracy similar to or better than previously published tools. Using CTAT-VIF, we systematically analyzed 33,763 tumor and normal samples, including over 1,000 cancer cell lines (**Supplemental Table S1**). This effort has contributed to expand and refine the existing catalog of biological samples with viral associations, genome insertions, and recompute and reassess viral insertion hotspots according to viral diversity and functional impact.

**Figure 1:**
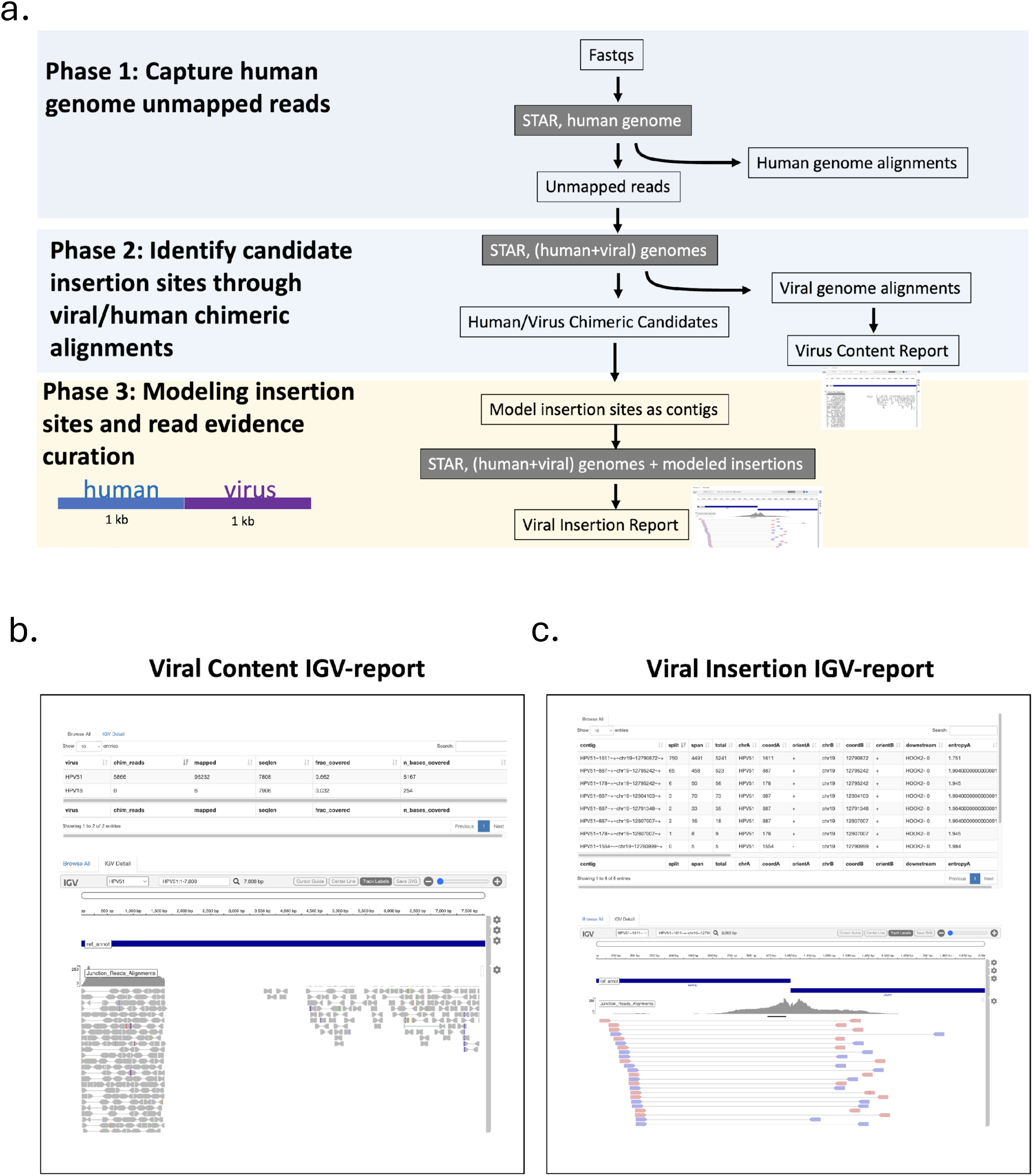
CTAT-VirusIntegrationFinder Workflow, Accuracy Assessment, and Visualizations. (a) The CTAT-VirusIntegrationFinder workflow involves three phases: (1) capturing human genome-unmapped reads, (2) identification of human::virus chimeric reads and virus-mapped reads from the genome-unmapped reads from phase-1, and (3) modeling virus insertion sites and evidence quantification. (b,c) Interactive IGV-report visualizations provided as outputs include (b) viral genome alignments yielding evidence for virus content and (c) read alignment support for modeled viral insertions derived from phase-3.

## Results

### CTAT-VirusIntegrationFinder Workflow

The CTAT-VIF computational workflow is a three-phase computational pipeline that harnesses the capabilities of the STAR aligner: (1) capture reads that either do not map or only partially map to the human genome; (2) for these partially mapped or unmapped reads, identify those that either fully map to viral genomes or as chimeric sequences between human and viral genomes; and (3) assess of virus insertion candidates through virus insertion modeling and the quantification of supporting evidence (**Figure 1a**). The viral genomes database used in phase-2 leverages a comprehensive collection of 962 human viruses, including reference viral genomes pertinent to oncogenic HPV, HBV, EBV, in addition to other viral genomes relevant to cancer or human disease (**Supplemental Table S2**).

### Virus Integration Accuracy Assessment and Benchmarking Comparison

To evaluate the accuracy of virus insertion detection across a diverse collection of viruses, we applied CTAT-VIF to simulated 50 base paired-end (PE) reads at either uniform (100x) or variable (1-20x) coverage levels. In these simulations, we introduced 100 random insertions per virus across 100 samples, with each of the 962 viruses randomly integrated in each sample.

Viral insertion candidates identified with at least 5 supporting reads in CTAT-VIF phase-2 were used for insertion contig modeling, realignment, and quantification. The final phase-3 achieved high recall (>99%) and high precision (>98%) across a range of 1 to 10 minimum chimeric evidence supporting reads (**Supplemental Figure S1a,b**). The CTAT-VIF phase-3 contribution to virus insertion detection accuracy over phase-2 was more pronounced with variable read coverage, where insertion modeling demonstrated a noticeable enhancement in quantifying virus evidence reads (**Supplemental Figure S1c**).

We further compared CTAT-VIF against the aforementioned published methods, focusing specifically on HPV insertion detection. To this end, we established test sets encompassing clinically relevant high-risk HPV strains, including HPV 16, 18, 31, 33, 35, 39, and 45. For each of these strains, we simulated 50 insertions per sample across triplicates (70 base PE reads at 50x coverage). All methods demonstrated high precision and recall (>90% on average); however, CTAT-VIF outperformed all other methods, emerging as the most accurate overall (**Figure 2**).

**Figure 2:**
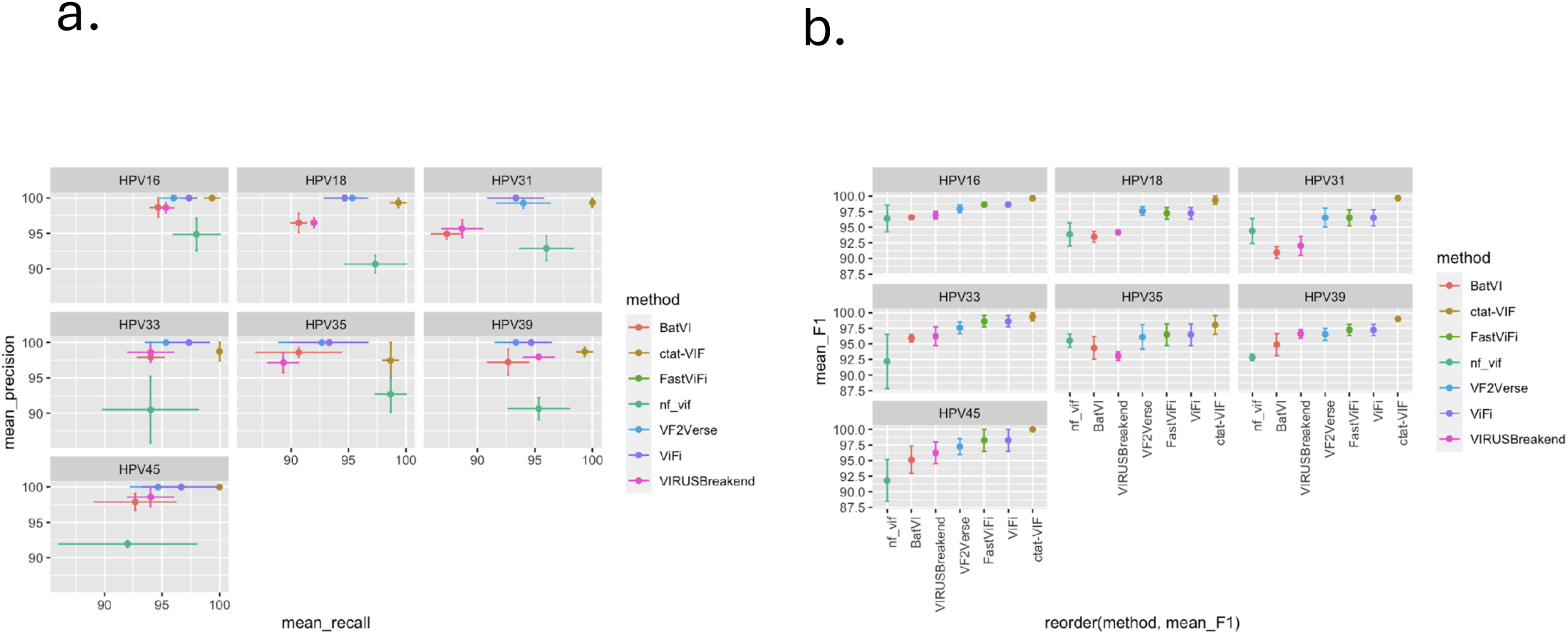
Benchmarking and comparing virus integration accuracy for related methods. Virus integration detection was evaluated for CTAT-VIF in comparison to six previously published methods using simulated data for seven pathogenic HPV strains with 50 insertions per virus in triplicates. per HPV strain. (a) Precision/recall and (b) F1 accuracy statistics according to HPV strain; error bars indicate standard error.

To evaluate CTAT-VIF performance using tumor-derived sequencing data, we analyzed previously published datasets comprising hybridization capture, whole-genome, and global transcriptome sequencing data derived from cervical cancers and associated cell lines as described in (Hu et al. 2015). The earlier-published work identified 3,666 HPV insertions in 103 out of 135 total hybrid capture (HYB) samples. Application of CTAT-VIF, however, unveiled a considerably larger number, detecting 17,928 HPV insertions in 125 out of 135 samples. Sub-selecting insertion regions at the most highly supported sites within 1kb genomic intervals and excluding a single sample (S12) that was unavailable in the SRA database among the other specimen in this collection revealed 2989 insertion sites from (Hu et al. 2015) and 9315 insertion loci identified by CTAT-VIF. When considering the most minimal supporting evidence of 1 supporting read alignment, only 935 of the previously published insertion sites were confirmed. Increasing the minimum read support threshold to 10 raised the agreement to nearly 75%, albeit with a drastic reduction in the total number of insertions (317 from CTAT-VIF and 238 from (Hu et al. 2015)). The degree of read support for the agreed-upon insertion loci was well-correlated (R=0.75), but the number of insertion loci per sample exhibited a lower correlation (R=0.41), with notable divergences evident (**Supplemental Figure S2a,b**). For instance, CTAT-VIF identified 742 HPV33 and 582 HPV11 insertion loci in HYB samples T2122 and T2116, respectively, whereas no insertions were previously documented for these samples.

Restricting our analysis to the subset of 175 Sanger-validated insertion sites, we observed similar rates of concordance with the previously detected insertion loci that exceeded minimal read support criteria. These 175 insertion sites can be categorized into 127 insertion loci binned at 1kb genomic intervals. Among these, CTAT-VIF identified 92 (72%) insertion loci that were validated by Sanger-sequencing insertion loci and revealed a high correlation among read support for shared loci (R=0.92**, Supplemental Figure S2c**).

### Virus Content Detected Across Diverse Tumor and Normal Tissues

Leveraging CTAT-VIF, we surveyed virus content and integrations across 33,762 samples involving 13,773 study participants or cell lines across the following distinct sources: The Cancer Genome Atlas (TCGA, (Cancer Genome Atlas Research et al. 2013)), the Genotype-Tissue Expression (GTEx, (Consortium 2013)) project, the Therapeutically Applicable Research to Generate Effective Treatments (TARGET) pediatric cancer project (Ma et al. 2018), the Cancer Cell Line Encyclopedia (CCLE, (Barretina et al. 2012)), and individual studies of HPV in cervical cancer (Hu et al. 2015; Kamal et al. 2021) and anal squamous cell carcinoma (Morel et al. 2019) (**Supplemental Table S1, Supplemental Figure S3**).

We initially restricted the detection of virus content across samples by imposing a requirement of a minimum of 10 viral genome-mapped reads covering at least 500 bases of the viral genome. Under these criteria, we captured evidence of a diverse array of viruses, with representation spanning all cohorts and sequencing modalities (**Supplemental Figure S4).** A primary concern arises due to the possibility that viruses detected within sequencing data could be attributed to contamination (e.g. cross-contamination where genetic material from a previous sample remains on equipment or surfaces; index hopping, when a DNA fragment from one library attaches to the barcode (index) of a different library during cluster generation on the flow cell). One notable sequencing contaminant was Ebola virus, detected in multiple GTEx samples, including several sourced from the same individuals (verified through personal communication, GTEx). Recognizing such sequencing contamination highlights its potential to impact the interpretation of the results, introducing confounding factors when considering potential biological associations, as seen in the case of HPV38 detected in uterine carcinoma.

Consistent with an earlier study ((Kazemian et al. 2015)), we detected HPV38 among TCGA uterine carcinoma (UCEC) samples; however, the levels of HPV38 were mostly at trace levels, detected in 46 UCEC RNA-seq samples, with only 2 samples slightly exceeding 10 RPM. In agreement with (Kazemian et al. 2015) we did not detect HPV38 among the corresponding whole exome sequencing (WXS data).

The consistent detection of virus within a biological sample, verified through orthogonal methods such as RNA-seq and WXS, greatly bolsters support for the detected biologically relevant virus content. This holds true except in cases where the presence is indicative of common trace-level contaminants. Indeed, within matched TCGA samples analyzed with multiple platforms, we found orthogonal validation of virus content via RNA-seq and WXS, revealing evidence of HPV in cervical and head and neck cancer, EBV in stomach cancer, and HBV in liver cancer (**Figure 3a**). In the context of these well-established and biologically relevant viral associations, the majority of detections manifest with at least 10 RPM in RNA-seq data, while matched WXS data display variable levels of supporting evidence. RNA-seq emerges as a more robust detection method for viruses that demonstrate consistent detection across both RNA-seq and WXS, particularly for HBV.

**Figure 3:**
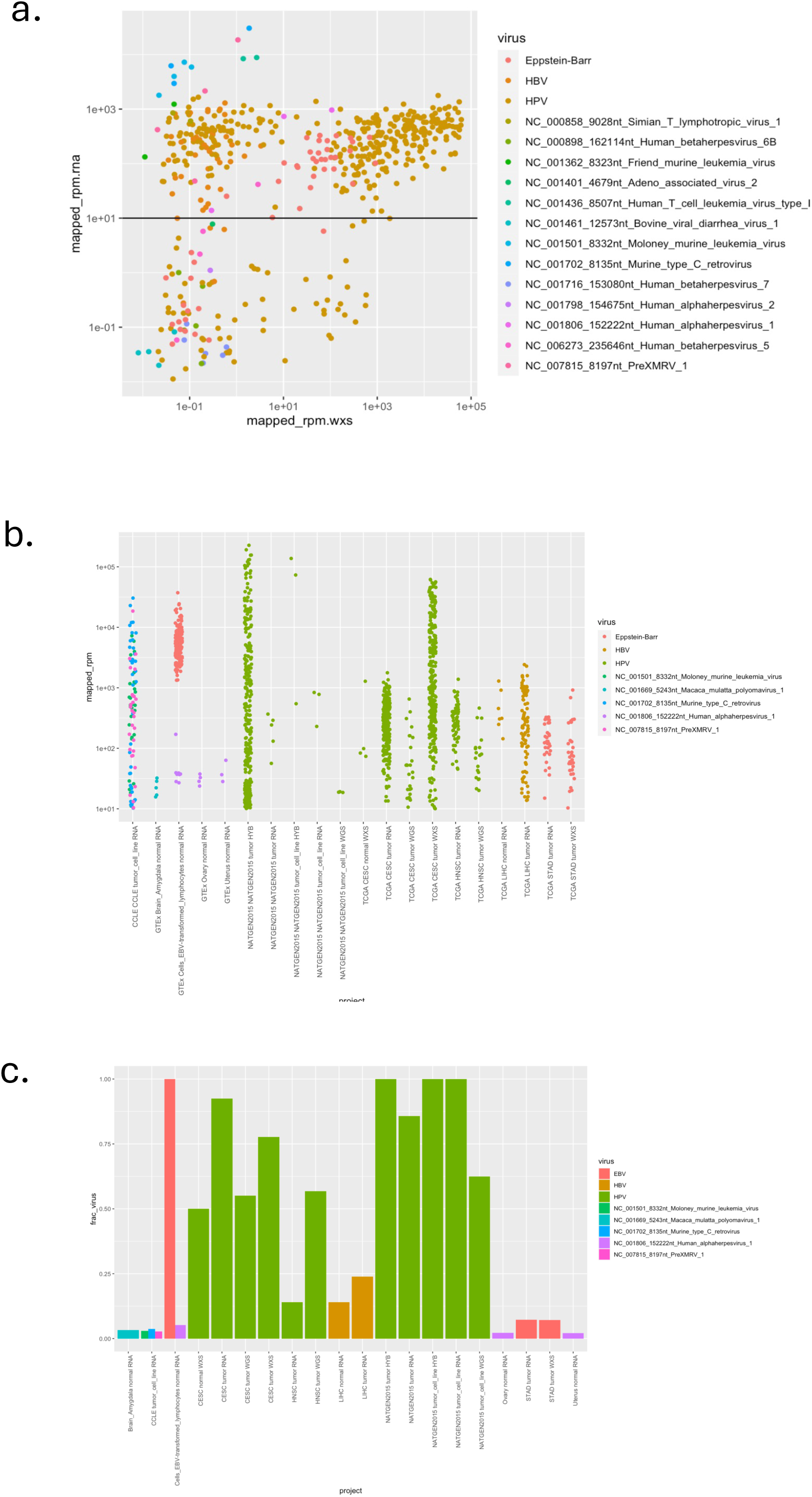
Virus content detected according to project cohort and sequencing modality. (a) Comparison of virus content and viral read quantifications according to RNA-seq and WXS for matched samples. For RNA-seq, most matched viral detections of pathogenic viruses with known disease associations (HPV, EBV, and HBV) are concentrated at least 10 rpm (black horizontal line), a threshold we define for discriminating trace-level virus detection. (b) Viral rpm observed according to sample type and sequencing modality above trace levels. (c) Fraction of project sample types found with corresponding virus detected above trace levels with sequencing modality indicated.

Upon implementing the more stringent 10 RPM threshold for virus content detection, we mostly observed associations consistent with the well-known virus and sample type relationship (**Figure 3b,c**). These include HPV in cervical (92%) and head and neck cancers (14%), HBV in liver cancer (24%) and tumor-matched normal liver samples (14%), EBV in stomach cancer (7%) and, perhaps unsurprisingly, EBV in all EBV-transformed lymphocyte samples from GTEx (100%). Among the cancer cell lines, approximately 5% (48/1018) showed evidence indicative of a common cell culture viral contaminant, the xenotropic murine leukemia viruses (MLVs, (Arias and Fan 2014)).

Beyond the prevailing patterns mentioned above, we found evidence of rare samples exhibiting notable virus content. Occurrences of HPV outside of cervical and head and neck cancers were infrequent. For instance, in the TCGA sarcoma tumor RNA sample SARC-MJ-A68H-TP, evidence of HPV51 presence was noted at 1.2k RPM, covering 72% of the HPV51 reference genome.

Furthermore, GTEx participant 1KXAM showed evidence of HPV182, a rare HPV strain related to HPV15 and HPV80, detected by CTAT-VIF on suprapubic sun-not-exposed skin at 386 RPM.

Herpes simplex virus 1 (HSV-1) is commonly detected at trace levels, and present in less than 2% of the various sample types surveyed. However, an exceptional case involving GTEx participant X4EP highlights a rare occurrence. In this sample, multiple brain tissues exhibited evidence of likely acute HSV-1 infection, with over a million HSV-1 reads detected (ranging from 19k to 162k RPM) across different brain regions, covering 96% of the HSV-1 genome. HSV-1 was sporadically found at high levels in other tumor and normal samples, including a normal thyroid sample (GTEX-1MCC2-0526-SM-DHXJS, 2.1k RPM), esophagus tumor (TCGA ESCA-LN-A8I0-TP, 1.3k RPM), stomach tumor (TCGA STAD-RD-A7BS-TP, 960 RPM), and head and neck tumor (TCGA HNSC-BA-5558-TP, 733 RPM).

The TARGET pediatric cancer samples were almost entirely devoid of virus content above trace levels. However, in the TARGET pediatric RT-tumor sample TARGET-52-PASILR, evidence of Parechovirus presence was detected at 31 RPM. Parechovirus is commonly associated with infections in children and is generally not harmful. Mastadenovirus C was detected at trace levels (<10 RPM) in several TARGET samples, while HSV-1 presence was not observed. Human immunodeficiency virus (HIV) was detected at levels below trace in 45 TARGET tumor or acute myeloid leukemia (AML) samples and was found in minimal quantities elsewhere.

### CTAT-VIF Integration Data Quality Assessment and Filtering Potential Contaminants

The number of identified virus insertions and their corresponding samples depends on the selected minimum evidence threshold (**Supplemental Figure S5a,b**). Requiring minimal support for an insertion identified by CTAT-VIF (1 evidence read from the final phase-3 insertion modeling and read realignments; see **Methods**), we identified a total of 37,184 virus insertions across 1323 samples. Increasing the minimum evidence to 2 reads yields approximately half the number of insertions (19,031) across three fourths (1083) of those samples. The decrease in sample count is mostly attributed to EBV insertions with minimal supporting evidence, while the prevalence of samples containing HPV or HBV insertions remain largely unchanged (**Supplementary Figure S5c,d**). Only a small fraction, 1.3% (with a minimum of 2 reads) of insertions were identified in samples that did not meet our previously established criteria of sufficient viral read content (minimum of 10 RPM and coverage of at least 500 viral genome bases). We categorized these corresponding insertion sites as suspicious and subsequently excluded them from any further analyses.

Cross-sample contamination, if present, could artificially detect common insertion sites across multiple samples. Alternatively, shared insertion sites may genuinely reflect true biological mechanisms, such as viral preference for inserting at specific genomic locations, or, in the case of transcriptome data, the use of common splice sites proximal to otherwise distinct genome insertion sites. Among the 37,184 instances of virus insertion across samples (involving 36,266 unique human genome sites), we observed 807 instances (involving 286 unique human genome sites) that were shared across samples from different sample donors. For example, a frequently shared insertion site involved an EBV spliced insertion detected in 22 GTEx EBV-transformed lymphocyte RNA-seq samples (EBV:NC_009334:40194 spliced into GRCh38:chr12:128814323). This site splices transcription from the EBV BHLF1 locus into the antisense direction of the SLC15A4 gene. Curiously, SLC15A is one of 10 Systemic Lupus Erythematosus (SLE) risk genes, suggesting potential biological relevance (Afrasiabi et al. 2022). In contrast, a commonly shared DNA insertion site involved an HPV16 insertion detected in 11 cervical tumor samples (HPV16:7267, chr3:60594700) at the FHIT locus, as identified in hybrid capture samples by (Hu et al. 2015), with several of these FHIT insertions labeled here as suspicious. Among all candidate insertions, only 88 (0.2% of the total insertions) had read evidence support at < 1% of the most highly supported shared insertion site, yielding little evidence for contamination.

For 252 RNA-seq samples containing high-confidence candidate insertions, we had matching WXS, WGS, or hybrid capture sequencing data available, providing orthogonal evidence for these insertions. Altogether, these samples contributed 6251 (17%) of all candidate insertions. Among them, approximately half (127) exhibited identical and matching human genome and viral genome breakpoints across the RNA-seq and the other sequencing approaches. However, not all orthogonally supported breakpoints could be validated at a single base resolution due to biological factors such as RNA-seq splicing versus direct genome integration coverage in DNA-seq, as well as bioinformatic variability in alignments at or near insertion breakpoints. Allowing up to 10kb between RNA-seq and other-seq insertion sites, we identified a total of 1189 concordant virus insertion sites in RNA-seq and other sequencing data for 220 of the 252 participants.

### Diverse viruses detected as genome insertions across tumor and normal tissues

We curated a data set of high-confidence insertions across tumor and normal tissues/cell lines, yielding 36,708 total insertions across tumor and normal tissues (**Supplemental Table S3**). Restricting counts of insertions to unique virus genome breakpoints, we identified 24,121 virus ‘break-ends’ (terminology adapted from (Cameron et al. 2021)). Grouping virus insertions within 10kb genomic intervals within samples refined our dataset to 29,164 insertion loci, encompassing 20,901 virus break-end loci. Among these, approximately ¾ of all virus insertions corresponded to HPV strains (77% insertion loci), followed by EBV (12%) and HBV (3%). Other viruses, such as HSV-1 and the MLVs, displayed single to multiple insertions per sample, with some samples containing up to hundreds of insertions.

Insertion counts and virus break-end counts were highly correlated (R=0.94, **Supplemental Figure S6a**). The number of insertion loci (grouping insertions within 10kb intervals) per sample range widely, from single insertions to over one thousand, depending on the experimental approach and virus type. Hybridization capture detected the highest insertion counts, whereas low coverage (∼6x) whole genome sequencing detected the lowest (**Supplemental Figure S6b**). Most samples contained a small number of viral insertions (median of 2 to 5) per sample (**Supplemental Table S4**). Samples with highest insertion loci counts were those from cervical tumors samples of the hybrid capture study conducted by (Hu et al., 2015). For instance, sample T2122 featured 2609 HPV33 virus insertion loci (999 virus break-ends), and sample T8729 presented 1518 HPV16 virus insertion loci (1215 virus break-ends). The counts of insertion loci (or break-end) were positively correlated among RNA-seq and matched WGS (R=0.86) or WXS (R=0.49), but not with hybrid capture (R= -0.3) (**Supplemental Figure S6c**).

Typically, less than half of DNA-seq detected insertions were recapitulated by RNA-seq, presumably due to insufficient or lack of transcript expression across corresponding sites (**Supplemental Figure S6d**). This discrepancy could also be attributed to the possibility of HPV insertions occurring in intergenic regions and not expressed, which would remain undetected in RNA-seq datasets (Yu et al. 2024).

Most samples exhibiting HPV or HBV content also exhibited evidence of corresponding viral insertions, with virus content and insertions detected across all sequencing modalities (**Figure 4, Supplemental Table S5**). While EBV content was consistently detected in 7% of stomach tumor samples (**Figure 3c**), insertions were only detected in the WXS samples; none were found in the RNA-seq samples. EBV insertions were detected in 98% of the EBV-transformed lymphocyte RNA-seq samples.

**Figure 4:**
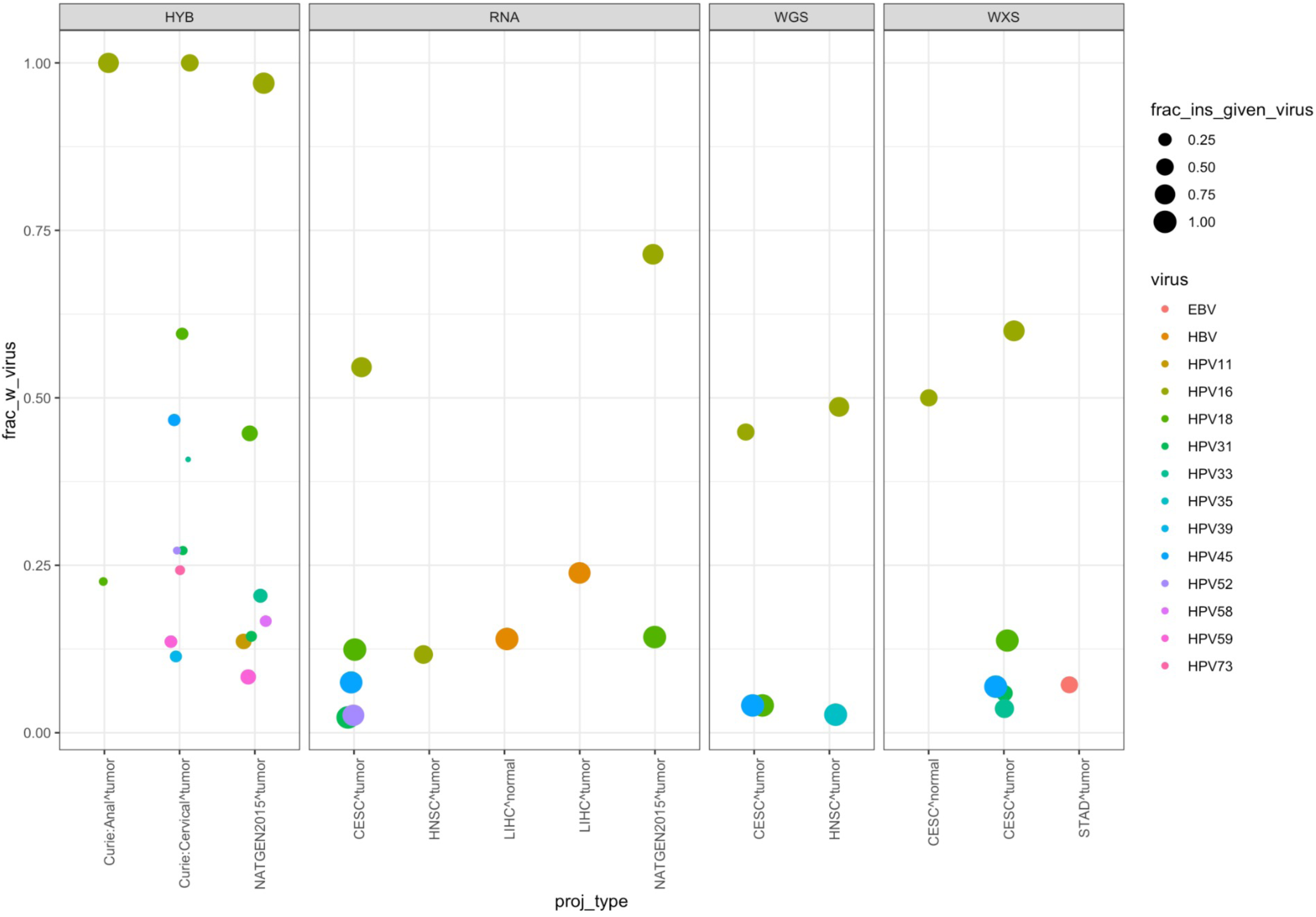
Virus insertion prevalence by virus, project cohort, and sequencing type. (a)Virus content and insertion prevalence by virus, project cohort, and sequencing type, restricted to those sample groups with a minimum of 2% of samples found with viral insertions (excluding ebv-transformed lymphocytes (not disease relevant) and the (Hu et al. 2015) non-hybrid-capture samples (too few samples)).

Rare examples of certain viral insertions found among surveyed samples included the following: A cell line from CCLE (BCP1_HAEMATOPOIETIC_AND_LYMPHOID_TISSUE) exhibited evidence of KSHV with 75 insertion loci (44 break-ends); A single TCGA liver tumor sample (LIHC-ZS-A9CF^LIHC) showed 3 insertion loci (3 break-ends) of Adeno_associated_virus_2; And another rare and peculiar example involved the aforementioned mentioned GTEx normal participant X4EP presenting evidence of 1878 HSV-1 insertions (281 loci) across six distinct brain areas: amygdala (819 insertions), anterior cingulate cortex (577 insertions), hypothalamus (331 insertions), and hippocampus (190 insertions). An additional insertion was found in both the cerebellum and the nucleus accumbens basal ganglia. Altogether, only 32 of these insertion sites (<2%) were shared across at least two brain regions within this individual.

### Functional impacts of insertions

Viral genome insertions are frequently linked to genome instability and altered expression of proximal genes (Jox et al. 1997; Zhao et al. 2016; McBride and Warburton 2017). Accordingly, we detected evidence of copy number amplifications or increased expression levels for genes located at viral insertion loci (**Figure 5**). HPV16 (p=7.3e-19) and HPV18 (p=1.8e-4), the two most prevalent HPV strains, were significantly associated with focal copy number amplifications at their insertion sites (Wilcoxon rank sum test with Benjamini Hochberg correction). For example, in the TCGA head and neck tumor sample HNSC-P3-A5QF-TP, an HPV16 insertion coincided with a focal genome amplification estimated at 49 copies (**Supplemental Figure S7**). Interestingly, while HPV (p=3.2e-20), HBV (p=2.3e-24), and EBV (p=4.5e-41) insertions were significantly associated with increased expression of proximal genes, only HPV demonstrated a significant association with copy number amplifications.

**Figure 5:**
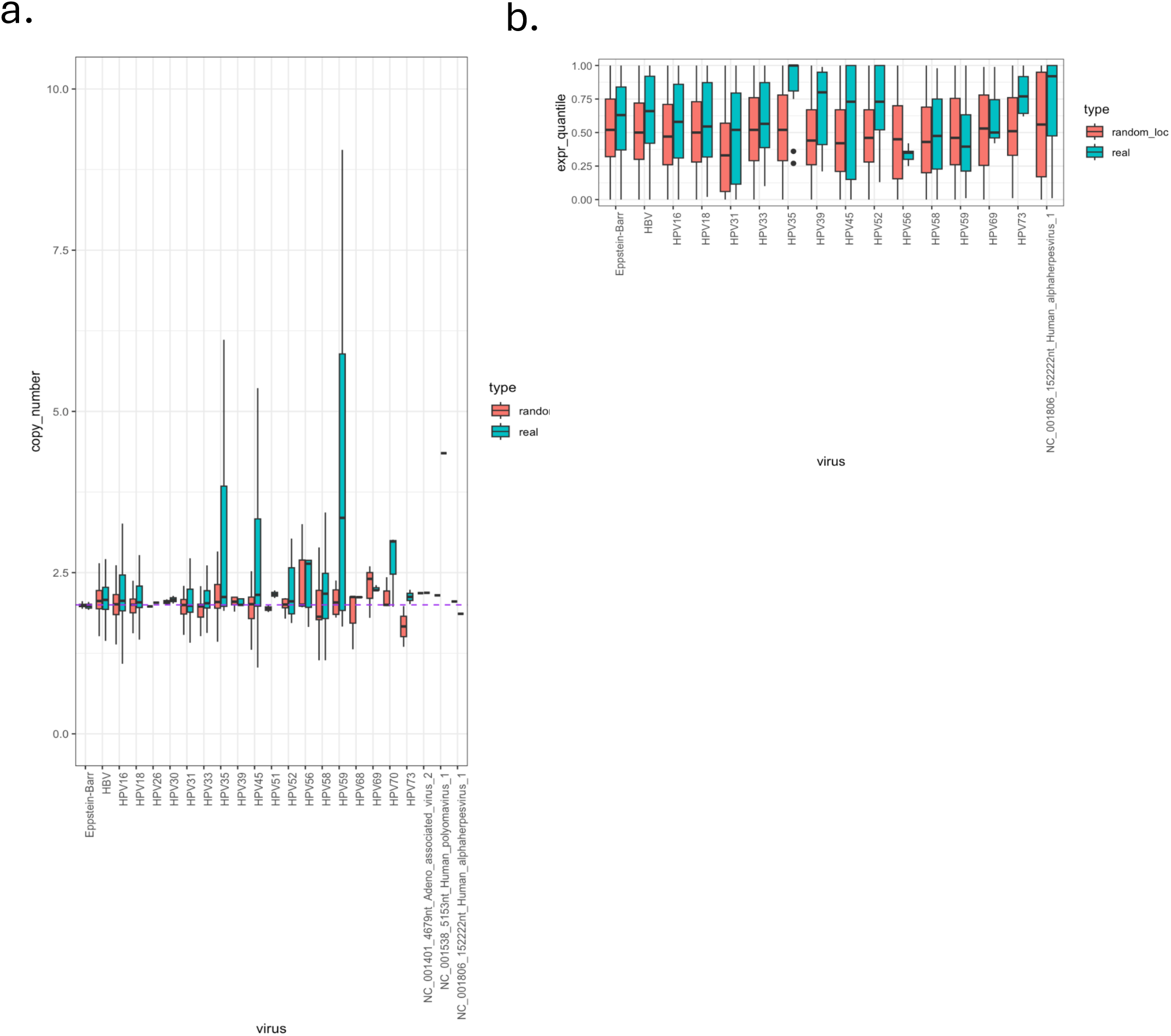
Functional impacts on expression and copy number aberration at viral insertion sites. (a) Distribution of maximum copy number detected within 10kb of insertion sites of corresponding virus types. (b) Distribution of expression value quantiles for aggregate gene expression within 10kb of corresponding viral insertion sites. For these analyses, 100 random sites were selected across the genome for each insertion within each sample to yield a null distribution. Statistical analysis leveraged the Wilcoxon Rank Sum Test comparing observed values to the corresponding null distribution.

Numerous fusion transcripts, both human/human or virus/human, were identified within 100kb of virus insertion sites. Among the 165 human/human fusion transcripts found proximal to viral insertion loci in 68 samples (**Supplemental Table S6**), several fusions were associated with known oncogenic events. Notably, we observed PVT1::MYC fusions in five cervical cancer samples (four with HPV16 and one with HPV35), as we previously noted in (Haas et al. 2023).

Additionally GRB7::ERBB2 fusions were found in three cervical cancers with HPV16 insertion. One cervical cancer (CESC-WL-A834-TP) contained a FGFR3::TACC3 fusion transcript mapping near a HPV16 insertion. Furthermore, the hallmark oncogenic fusion TMPRSS2::ERG characteristic of prostate cancer, was identified proximal to a MLV insertion in the Vertebral-Cancer of the Prostate (VCaP) prostate cancer cell line. Although the TMPRSS2::ERG fusion is a feature of the prostate cancer from which the VCaP cell line originated, the MLV insertion is presumably unrelated and attributed to a MLV cell culture contaminant.

Collectively, we identified 301 distinct human genes exhibiting chimeric splicing with viral transcripts across a total of 229 samples (**Supplemental Table S7**). This analysis utilized human reference exon splice sites in conjunction with known or cryptic viral splice sites. Within these virus/human fusion transcripts, several human genes are recognized as oncogenes (ERBB2, CD274(PD-L1), SRC, TERT, GRB7) or tumor suppressors (TP53, TP73, RAD51B, KMT2B, PTPN13).

The viruses predominantly involved in these chimerically spliced fusions include HPV (59%), EBV (15%), and HBV (11%). HPV spliced fusions primarily derive from known viral splice sites of early genes (E6, E7, E4, and E2). In contrast, EBV and HBV spliced fusions mostly derive from cryptic splice sites in each viral genome, as they do not consistently align with reference transcript annotations.

### Insertion Hotspots

HPV, HBV, and EBV were previously found to frequently insert at certain genomic loci referred to as hotspots (Hu et al. 2015; Bodelon et al. 2016; Zhao et al. 2016; Li et al. 2018; Kamal et al. 2021; Bi et al. 2023). Earlier identification of these hotspots relied on smaller datasets, used larger window sizes to aggregate insertions into hotspot regions, or lacked the rigorous filtering for potential artifacts and contaminants that we have implemented in our study. Leveraging our extensive collection of virus insertion sites, we redefined genomic insertion hotspots with higher resolution and accuracy. We examined the number of participants with virus insertions within 100 kb windows across the genome and captured those windows containing peak numbers of participants with virus insertions (ie. hotspot sizes) as candidate hotspots. Given the large number of insertion sites, it was anticipated and indeed observed that some candidate hotspots characterized by multiple virus insertions would emerge by chance, with distributions of random insertion hotspot sizes dependent on window size. To examine the occurrence of hotspots by chance (null model), we performed 10,000 trials by first randomizing the virus insertion positions across the genome and subsequently identifying hotspots (**Supplemental Figure S8a**). Notably, hotspots representing 6 or fewer participants were readily observed by chance, whereas those involving 9 or more participants were rare or not observed. By defining hotspots as those involving insertions from at least 9 participants, we identify 227 hotspots that represent roughly 1/4 of the virus insertions (10,145 / 36,708) (**Figure 6a**).

**Figure 6:**
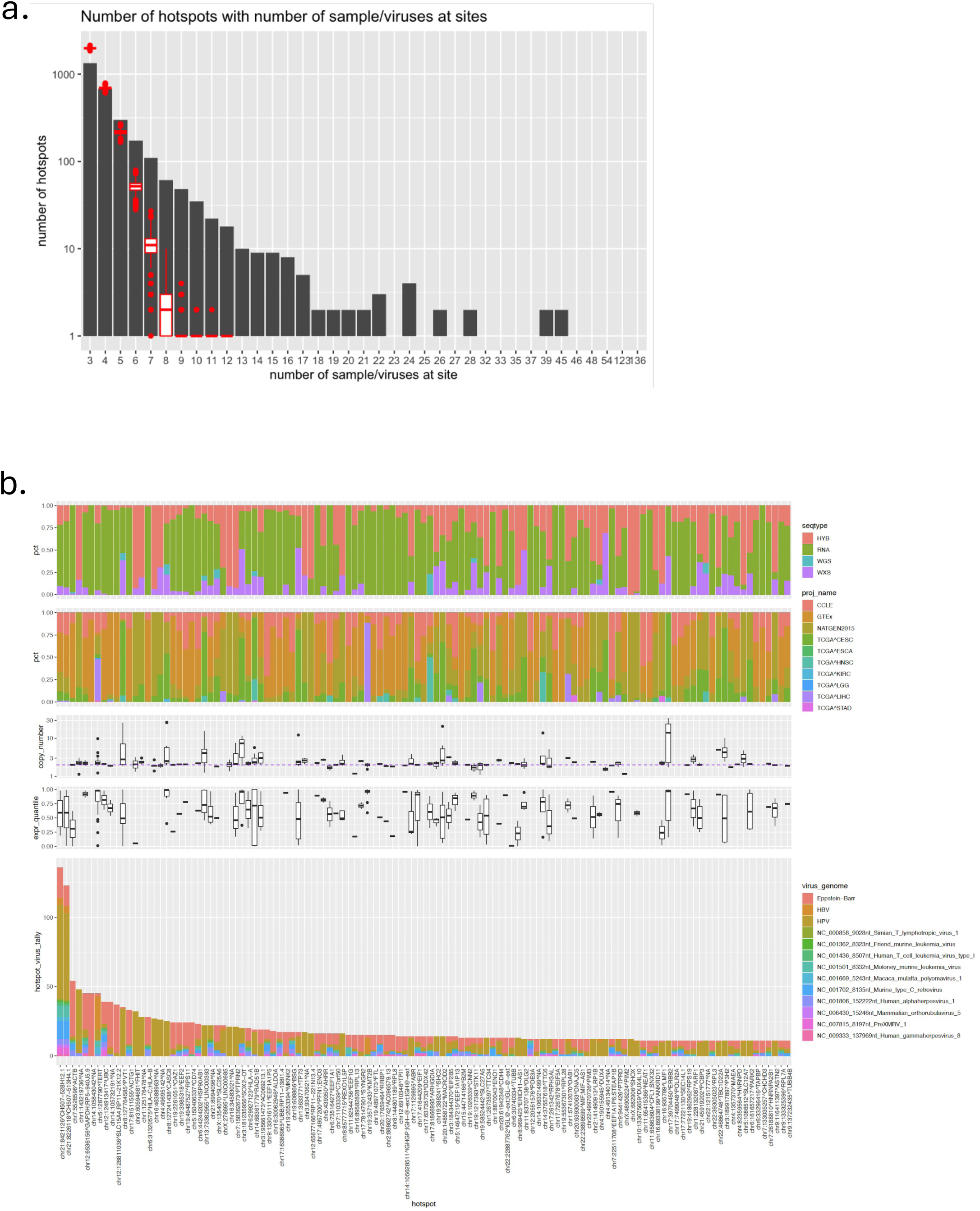
Virus insertion hotspots and characteristics. (a) Counts of hotspots observed according to sample representation using a window size of 100k. The distribution of hotspot sizes for random data using the same window size is shown in red and matches that in **Supplemental Figure S8a**. (b) Top viral insertion hotspots (min. 9 samples) observed and ranked according to sample representation (descendingly from left to right) with the following characteristics (top to bottom): sequencing type representation, project representation, distribution of maximal copy numbers observed within 10kb of insertion sites, aggregate expression quantiles for genes within 10kb of insertions, and counts of samples with corresponding hotspot insertions according to virus type.

Representation of top hotspots according to virus type, project cohort, sequencing data type, and potential functional impact - as ascertained via expression quantification (available for TCGA and GTEx) and copy number aberration (restricted to data availability in TCGA) - is shown in **Figure 6b**. Many of the hotspots appeared to be shared across different viruses, while others were specific to certain virus-type. Among the top hotspots, the two most prominent were localized to a position at approximately 8 Mb on GRCh38 chromosome 21, corresponded to insertions at the rDNA locus, and likely represents virus insertions within the expansive 45S rDNA array on chromosome 21. Other top hotspots included the housekeeping genes ACTB and GAPDH, which were affected by diverse virus types. When examining the functional impacts on expression at these hotspots, the predominant trend was an increase in expression levels, as observed by the distribution among the highest expression quantiles, with few exceptions (e.g., FHIT as described below).

Nearly half of the identified top hotspots exhibited HPV-specific enrichment. The top gene-associated insertion loci were located on chromosome 8, and they were associated with the oncogenes PVT1 or CASC8. This genomic region spans approximately1 Mb and includes a third defined hotspot at CASC11. These three chromosome 8 proximal hotspots collectively harbor insertions associated with MYC, CASC19, and CCAT1 (**Figure 7a**). Together, these hotspots were identified with insertions from 13% of the cervical cancers surveyed. Notably, this region also showed enrichment of copy number amplifications and relatively high expression levels, particularly at the CASC8 hotspot (**Figures 6b,7a**). Additional HPV-enriched hotspots were observed in regions characterized by the highest copy amplifications and expression levels.

**Figure 7.**
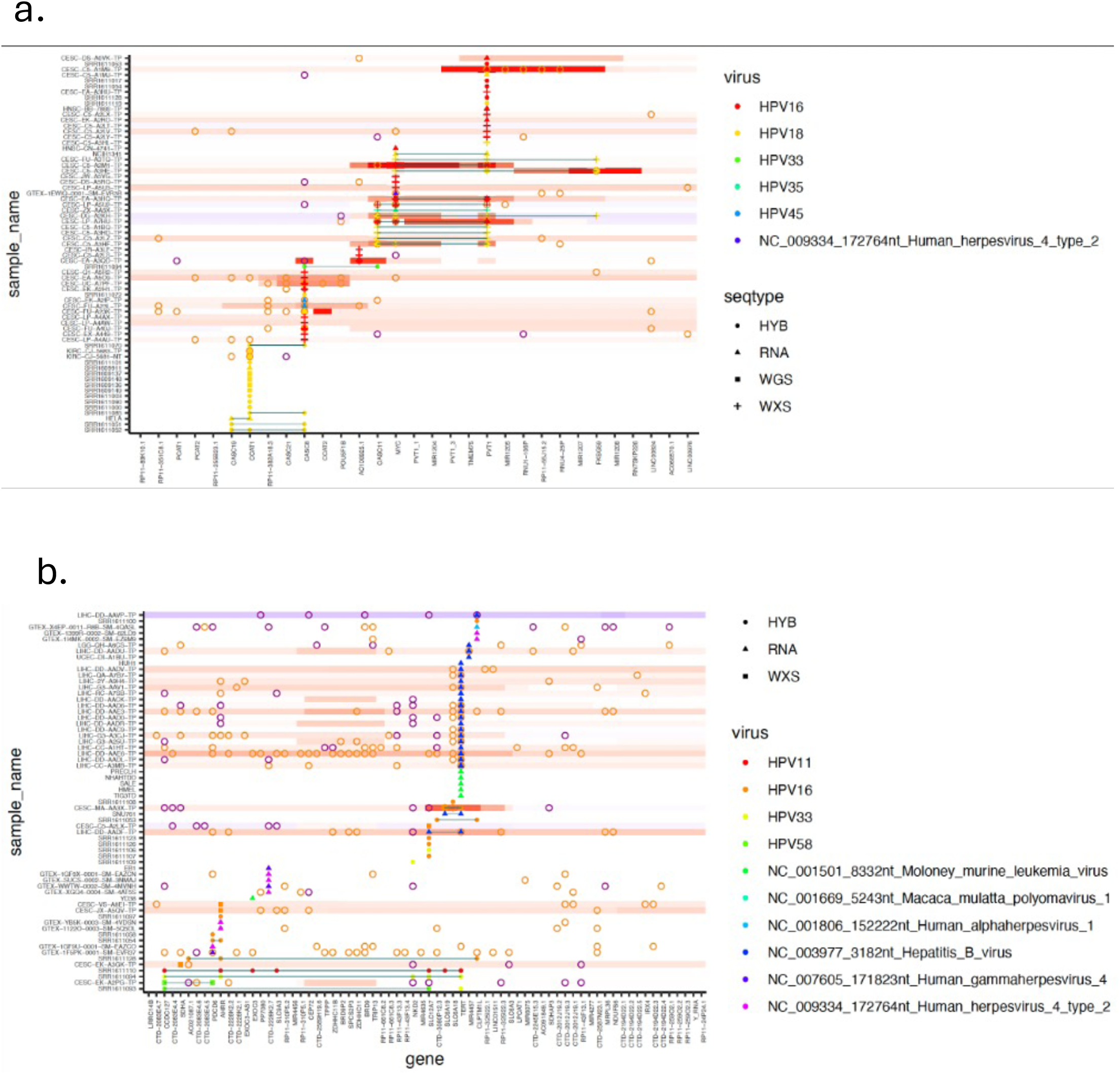
Insertions and functional impacts for top hotspots. (a) HPV insertion hotspot on chr8 involving PVT1, MYC, CASC8, and CASC11. (b) HBV insertion hotspot at the TERT locus. Insertion sites are shown according to most proximal gene (x-axis), sample (y-axis), virus type (color) and sequencing type (shape). Functional impacts are illustrated as follows: genes with expression values found in the top (>=0.95, magenta hollow circle) or bottom (<= 0.05 quantile, orange hollow circle) expression quantiles, and copy number aberrations as colored bars, involving amplification (red) or deletion (blue). Expression data is limited to TCGA and GTEx and copy number data are limited to TCGA.

These included LINC00393, PTPRN2, SOX2-OT, RAD51B, TP73, ABR, MACROD2, ERBB2, and P3H2. The prevalence of cervical cancer insertions at these hotspots varied across different project cohorts and sequencing types (**Supplementary Figure S8b**). For example, the HPV hotspot at the FHIT locus was detected in 22% of (Hu et al. 2015) cervical cancers surveyed by hybrid capture sequencing, whereas its detection was <1% via TCGA WXS data (**Supplemental Figure S8c**). Evidence suggests several of the FHIT insertions in the (Hu et al. 2015) data likely derived from sequencing cross-sample contamination hence inflating the frequency of this insertion hotspot (**Supplemental Figure S9a**). Furthermore, we identified two TCGA CESC samples (CESC-ZJ-A8QR-TP withHPV16 and CESC-4J-AA1J-TP with HPV18) in which FHIT gene insertions were evident from WXS but not RNA-seq. These samples displayed the lowest expression quantiles coupled with copy number deletions and hemizygous viral insertions (**Supplementary Figure S9b**), supporting that the FHIT insertions in these samples are genuine. Another hotspot involving LINC00393 was consistently detected in 4% of cervical cancers across both TCGA CESC and (Hu et al. 2015) cohorts. Several of these HPV hotspots also involved chimerically spliced fusion transcripts, either virus/human or human/human genes. Notably, fusion transcripts such as PVT1::MYC, CCAT1::PCAT2, ETV6::MACROD2, and GRB7::ERBB2 were identified, with the GRB7 and ERBB2 each generating fusion transcripts with additional partners at the ERBB2 hotspot.

HBV insertions were found particularly enriched at oncogenes TERT (in 5% liver tumors) (**Figure 7b**) and KMT2B (3% liver tumors), as well as at FN1 (4% tumor-matched normal livers). These loci are recognized as hotspots for HBV insertions (Zhao et al. 2016). Insertions at TERT and KMT2B hotspots were both associated with significantly higher expression levels of the respective genes, although no corresponding copy number changes were observed. Conversely, FN1 insertions were found in matched normal liver tissues rather than in the tumors, consistent with earlier reports (Ding et al. 2012; Shiraishi et al. 2014; Yoo et al. 2017; Budzinska et al. 2018; Furuta et al. 2018; Jang et al. 2021).

Nearly all (94%) of EBV insertions were detected in EBV-transformed lymphocytes within GTEx. These insertions frequently occurred at housekeeping genes ACTB and GAPDH, as well as the lymphocyte marker CD74, each associated with high expression of neighboring genes. Most of the top viral insertion hotspots identified in tumors displayed evidence of proximal EBV insertions originating from EBV-transformed lymphocytes within GTEx samples, or from HSV-1 insertions from GTEx participant X4EP infected brain samples.

We identified 432 samples with evidence of insertions at multiple hotspots. For example, sample SRR1611094 of (Hu et al. 2015) dataset had evidence of 127 hotspot insertions involving HPV33. The prostate cancer cell line VCaP had 68 hotspot insertions involving Murine retrovirus type-C, and GTEx EBV-transformed lymphocyte sample GTEX-1F5PK-0001-SM-EVR37 had 52 EBV hotspot insertions. In addition to insertions at multiple hotspots, some samples also displayed fusion transcripts linked to these insertions. For example, the cervical cancer sample CESC-C5-A2LX-TP had 18 HPV16 hotspot insertions along with fusion transcripts ERBB2::IKZF3, DCST1::ASPSCR1, and a virus fusion transcript involving HPV16-spliced with CIRBP.

## Discussion

The comprehensive characterization of cancer genomes and transcriptomes, crucial for understanding drivers of tumorigenesis, requires a robust suite of advanced analyses and tools. This toolkit must effectively handle complex analyses and tasks such as dissecting genomic structural and numerical aberrations, identifying mutations, deciphering fusion transcripts, and detecting oncogenic viruses. For studying oncogenic viruses, specialized tools should accurately establish virus identity, identify expressed viral genes, pinpoint accurate positions of virus insertions within the genome, and provide evidence for virus/human gene spliced transcripts generated at insertion loci. Here, we developed CTAT-VIF as part of our Cancer Transcriptome Analysis Toolkit, providing a rigorous approach to studying oncogenic viruses in cancer genomes and transcriptomes. Leveraging simulations with diverse virus-type genome insertions, we validated CTAT-VIF’s performance and demonstrated its high accuracy compared to previously published methods. Furthermore, we achieved a high validation rate based on earlier Sanger-sequenced insertions from (Hu et al. 2015). To build a more comprehensive catalog of human virus, sample type associations, genome insertions, and hotspots, we applied CTAT-VIF to over 30,000 samples across TCGA, GTEx, TARGET, CCLE, and sequencing data from previous published surveys of cervical and anal cancer. This comprehensive dataset included a variety of sequencing approaches, mostly RNA-seq, but also incorporated hybrid capture, WGS, and WES data where available and relevant for specific samples. Altogether, we identified 36,708 insertions and 227 hotspots (aggregating 10,145 insertions), mostly involving HPV in cervical and head and neck cancers, HBV in liver cancer, EBV in stomach cancer and EBV-transformed lymphocytes, as well as XMRV or related viral contaminants in cancer cell lines.

Several of the viral insertion hotspots were specifically associated with HPV or exhibited a strong association with HPV. These hotspots were frequently associated with focal copy number amplifications and increased gene expression at the insertion loci. Notably, insertions at these hotspot loci were often associated with known oncogenic human fusion transcripts or virus::human spliced fusion transcripts, well exemplified by the hotspot on chromosome 8 involving MYC, PVT1, and CASC8.

HBV was detected in liver cancer RNA-seq data, with prominent insertion hotspots limited to TERT and KMT2B in liver cancer, and at FN1 in tumor-matched normal liver tissues. In TCGA WXS data, the sensitivity for detecting HBV reads was ∼100 fold lower, and perhaps unsurprisingly, we were unable to identify HBV insertions from WXS data. While HBV insertions overall were significantly associated with increased expression of neighboring loci, no significant association with copy number alterations was observed.

Most of the EBV insertion hotspots were colocalized with insertions from other virus types in various samples including HPV, XMRV, and SV40. In addition, we detected a high prevalence of HSV-1 infections in brain samples surveyed from one of the GTEx participants, with insertions coinciding with EBV insertion hotspots across several of these samples. However, no significant association of EBV insertion with altered expression or copy number aberrations was observed. The viral diversity at EBV insertion hotspots, coupled with the limited functional impact on gene expression or copy number alterations observed at these sites, suggests that the detected EBV insertion sites may reflect regions of genome instability or regions otherwise more tolerant for integration of foreign DNA and perhaps less relevant to tumorigenesis.

In addition to copy number aberrations, most notably focal copy number amplifications, and increased expression of genes at insertion loci, the presence of oncogenic fusion transcripts found at these loci such as those involving PVT1, MYC, or ERBB2, are likely contributing factors to pathogenesis. The roles played by virus::human fusion transcripts originating from insertions remains unclear but could potentially contribute to pathogenesis and aid in neoantigen discovery to facilitate targeted immunotherapy efforts.

Virus detection can be confounded by sequencing contamination, as previously reported for HPV38 in uterine cancer and as we observed for Ebola in a small subset of the GTEx samples. For viruses with established biological significance, such as HPV, HBV, and EBV, viral reads mostly exceeded 10 rpm and in extreme cases over 100 rpm for non-hybrid-capture methods. Commonly encountered viruses found at trace levels, such as Herpesvirus and Mastadenovirus C, were mostly found below 10 rpm.

We found minimal evidence suggesting contamination that could interfere with the interpretation of virus integration. Instances where virus insertion sites featured identical breakpoints in both the virus and the human genome across samples from different participants, indicating direct physical virus/human genome breakpoint (and not involving spliced virus::human fusion transcripts), would raise significant suspicion. Earlier published results from the (Hu et al. 2015) study were earlier called into question for false positives and evidence of sequencing contaminants (Dyer et al. 2016; Hu et al. 2016). We also identified potential sequencing contamination involving identical insertions at the FHIT hotspot, which might overestimate the prevalence of FHIT insertions. Additionally, we identified two cervical cancer cases with evidence of HPV insertions at the FHIT locus, which led to a similar impact on reduced transcription at the FHIT locus coupled with copy number deletions (apparent hemizygous viral insertions). Insertions were additionally detected in separately sequenced cervical tumor samples (**Supplementary Table S3**), providing further support for the importance of the FHIT locus as an HPV hotspot with functional effect.

Applying CTAT-VIF to extensive collections of both tumor and normal samples across different sequencing modalities showcased its capabilities for detecting virus content and virus integration on a large scale. This application not only reaffirmed known virus and sample associations but also substantially enriched the catalog of genome insertion sites and hotspots. Pediatric and normal samples were found largely devoid of virus insertions, with few exceptions – HBV in normal liver, a few other interesting virus insertions, and special HSV-1 example in normal brain.

Cell lines serve as valuable models for studying insertions. While XMRV insertions were prevalent in cell lines, a subset of insertions showed biological relevance. Notably, HeLa cells displayed a chromosome 8 hotspot insertion with consequential effects including human::human and virus::human fusion transcripts. Several other biologically relevant insertions were identified. Certain cell lines offer potential for further in-depth exploration into the roles of virus insertions and their phenotypic effects. Furthermore, understanding the influence of virus insertions and any expressed viral products should be considered when studying these cell lines in targeted assays (eg. anticancer drug discovery).

While our CTAT primarily focused on exploring cancer biology through transcriptome studies via RNA-seq, the CTAT-VIF module has also demonstrated remarkable effectiveness in analyzing DNA-seq data from WGS, WXS, and hybrid capture sequencing data. Our analysis of WXS, WGS, or hybrid capture data provided orthogonal support for RNA-seq-defined insertion sites in samples. However, each data type has its own unique strengths and limitations for virus detection. RNA-seq proved particularly useful as it can detect both genome insertion sites and virus::human fusion transcripts, but requires active transcription in the regions of interest.

DNA-seq, especially hybrid capture sequencing, excels in surveying insertion sites across the genome but may not define virus::human spliced products or those insertions with the most functional relevance for understanding impacts of insertion sites. DNA-seq is crucial for capturing insertions that result in expression knock-outs or gene deletions, such as observed in the FHIT gene. The combination of hybrid capture and RNA-seq should prove most useful for more comprehensive characterization of viral integrations. Including HPV, HBV, and EBV capture as standard in exome capture kits would be beneficial for clinical assays, as WES becomes increasingly routine for tumor sequencing.

The interactive visualizations generated using IGV-reports simplify analysis of virus genome content and evaluation of the read alignment evidence supporting any reported genome insertions. Through concerted execution of CTAT modules, cancer transcriptomes can be more thoroughly evaluated for potential drivers of tumorigenesis, consider related events including the forms of fusion transcripts generated at viral insertion loci, and explore potential opportunities for future efforts in combating cancer in the new era of personalized medicine. Our open source CTAT-VIF software is freely available at https://github.com/broadinstitute/CTAT-VirusIntegrationFinder.

## Methods

### CTAT-VirusIntegrationFinder (CTAT-VIF) method and implementation

CTAT-VIF operates through three phases (**Figure 1a**): (Phase-1) captures reads that do not map concordantly to the human reference genome; (Phase-2), from these reads, those that map concordantly to viral genomes or align as chimeric human::viral sequences are identified; (Phase-3) involves modeling viral insertion sites in the human genome and quantification of corresponding read support. The entire workflow was implemented in the Workflow Description Language (WDL). These three phases are detailed below.

### CTAT-VIF Phase-1: capturing human genome un-mapped or partially-mapped reads

Reads were mapped to the human reference genome (GRCh38) using the STAR aligner with chimeric read alignments disabled (Dobin et al. 2013). Reads reported by STAR as unmapped were trimmed of low-quality bases using Trimmomatic (Bolger et al. 2014) followed by stripping of poly-A from read sequence termini. Reads at least 30 bases in length were retained and subject to Phase-2.

### CTAT-VIF Phase-2: identification of viral and human::viral chimeric reads

STAR was executed again using the reads captured from Phase-1 (genome-unmapped reads, cleaned and trimmed), searching a combined genome database incorporating the human reference genome with all 962 viral genome sequences, and allowing for chimeric alignments. STAR-reported human::virus chimeric read alignments were filtered to remove duplicates and those reads multi-mapping at >50 locations in the genome. Reads partially aligning to the mitochondrial genome were excluded. Chimeric events are defined according to human and virus breakpoint coordinates with supporting reads aggregated accordingly as evidence support. Chimeric events are ranked first according to alignment evidence type, with breakpoint defining split reads prioritized over paired-end spanning read pairs that lack precise breakpoint definitions, followed by prioritizing events with highest read support. Lower-ranking events within 500 bases of each the human genome and viral genome breakpoints of higher-ranking events are subsumed as supplementary evidence for higher ranking events; primarily intended to group spanning chimeric read pairs with breakpoint-defining split reads as individual events.

Chimeric read alignments were further grouped according to virus breakpoint (breakend) and multi-mapping reads were greedily assigned to chimeric events with shared virus breakends having highest cumulative reads assigned. Highest supported chimeric events for each virus breakend were assigned as primary and remaining events were assigned as non-primary. Non-primary events with less than 10% read support of primary events for the same virus break-ends or having more than 25% of alignments as multimapping were excluded. Finally, any events with fewer than 5 supporting read alignments were excluded.

Events defined as primary were defined as insertion candidates for modeling and reassessment in Phase-3. These primary events were further supplemented with highest-supported non-primary shared virus break-end events having unique insertion flanking sequence on the human genome (32 bases) and considered likely independent insertion events. These selected candidates are treated as representative of all non-primary insertion candidates with shared virus break-ends and flanking human genome sequences.

Those non-chimeric and otherwise concordant read alignments to viral genomes were separately leveraged to define evidence of virus content.

### CTAT-VIF Phase 3: virus insertion modeling

For each insertion candidate, an insertion event is modeled fusing 1kb each of human genome sequence and viral genome sequence centered at the breakpoint. The unmapped reads from Phase-1 were realigned to the combined human reference genome, the viral genome database, and the new database of insertion-modeled contigs with chimeric alignments disabled, focused on capturing those reads that best align to the virus insertion contig models instead of the reference human genome or viral genomes. The alignments to insertion contigs were analyzed and filtered as follows, requiring non-duplicate alignments, mapping quality > 0, alignment percent identity >= 95%, read sequence Shannon entropy >= 0.75, soft- or hard-clipping < 10 bases, and at least 30 bases aligned on both sides of the insertion breakpoint. Alignments passing these filtering criteria were tallied and assigned as read quantifications for corresponding insertions. Non-primary Phase-2 candidates with shared virus break-ends and flanking human insertion sequences with Phase-3 surviving representatives were recovered, assigned imputed quantification values by proxy, and reported in the final phase-3 insertion predictions.

### Application of CTAT-VIF to available tumor and normal samples

CTAT-VIF v1.5.0 was run on TCGA v11, GTEx v8, CCLE, TARGET, and (Hu et al. 2015) samples using Terra. Cervical and anal cancer samples from (Morel et al. 2019; Kamal et al. 2021) were separately processed separately leveraging the computational infrastructure at the Curie Institute. All samples processed are described in **Supplementary Table S1**.

## Resource availability

Further information and requests for resources or code should be directed to lead contact Brian Haas.

## Materials availability

This study did not generate new unique reagents.

## Key resources table

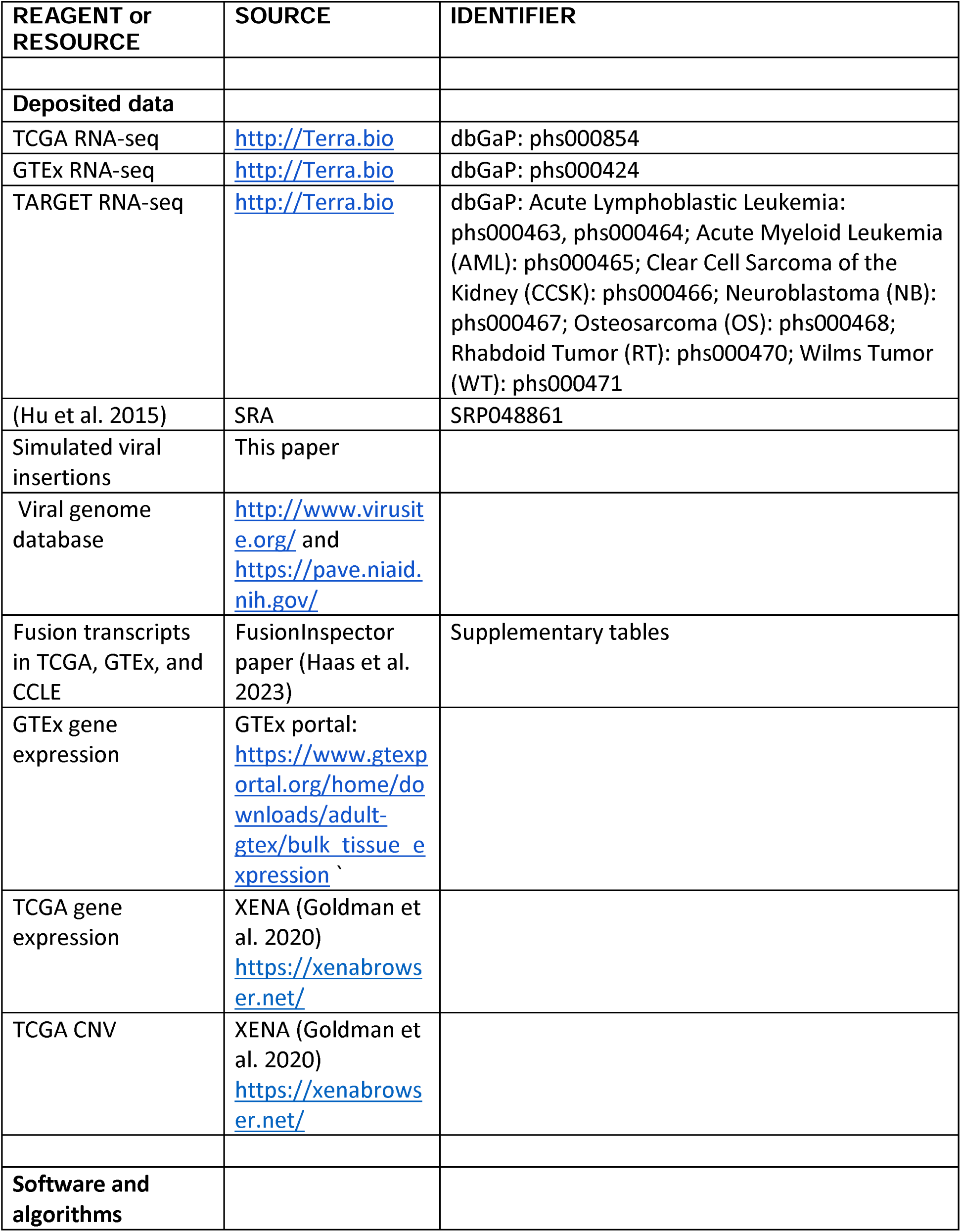
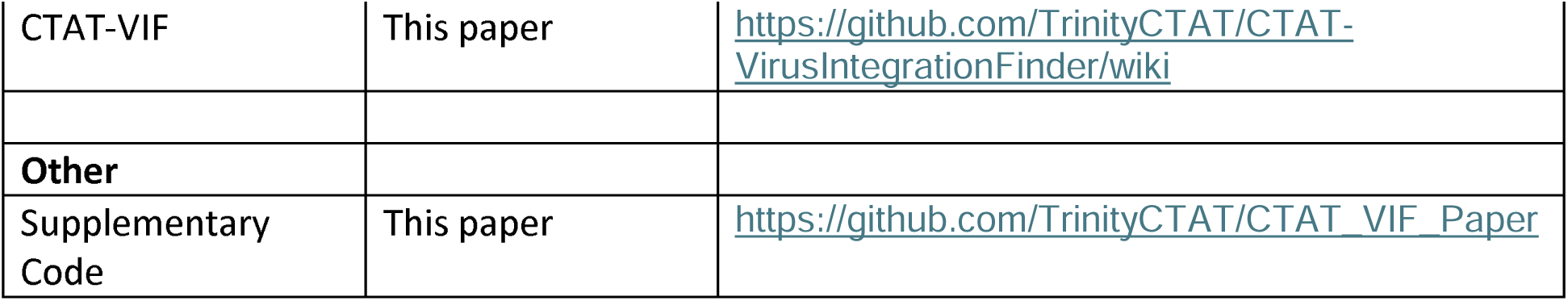

