## Supplemental Figures for "Pathogenic Viruses, Genome Integrations, and Viral::Human Chimeric Transcripts Detected by VirusIntegrationFinder Across >30k Human Tumor and Normal Samples"

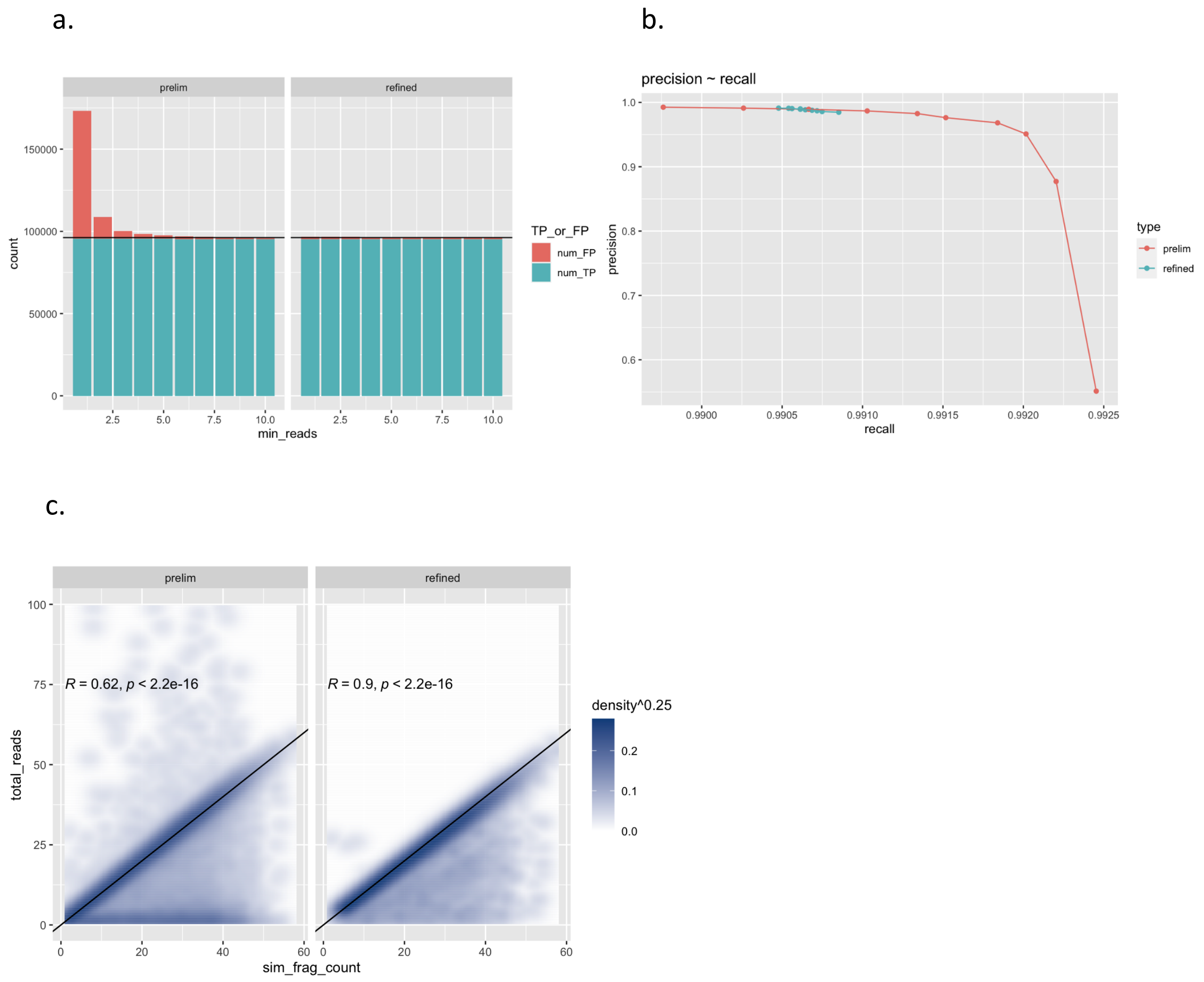


**Supplemental Figure S1: Comparison of evidence quantification agreement for CTAT-VIF phase-2 (prelim) vs. phase-3 (refined) with simulated insertions.** (a) Numbers of true (TP) or false positives (FP) according to minimum evidence read count threshold from 1 to 10 chimeric reads at 100x simulated uniform sequencing coverage. (b) Precision and recall values according to minimum evidence read thresholds in (a). Precision escalates from 55% at 1 chimeric evidence read to >97% when 5 chimeric evidence reads were imposed. Further refinement of phase-2 (prelim) candidates occurred in phase-3 (refined) through the application of a minimum of 5 chimeric evidence reads and virus insertion modeling and realignments. This iterative process retained high recall (>99%) and high precision (>98%) across the range of 1 to 10 minimum chimeric evidence supporting reads. (c) Phase-3 contribution to virus insertion detection accuracy over Phase-2 was more pronounced with variable read coverage, where insertion modeling demonstrated a noticeable enhancement in quantifying virus evidence reads. Similar results were obtained with 100 or 150 base PE reads (**Supplementary Code**).

a.

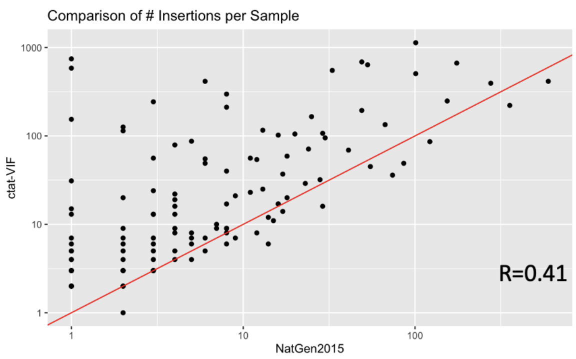


b.

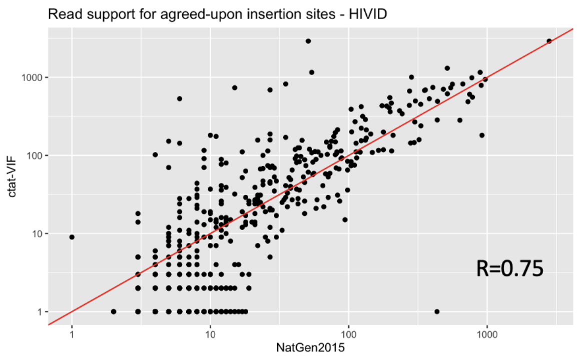


c.

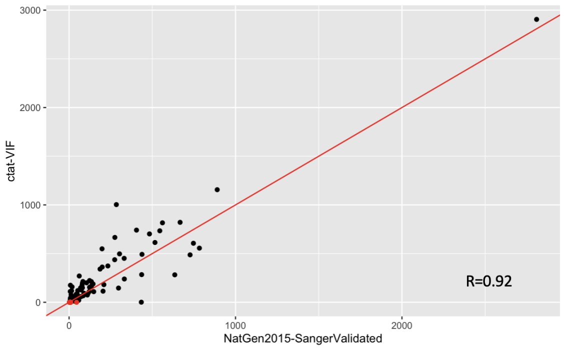


**Supplemental Figure S2: Comparison of virus insertion prevalence and read support between CTAT-VIF and previously reported analysis of cervical cancer cohort Hu et al., 2015 (NATGEN2015).** (a) Comparison of numbers of insertions per sample, (b) quantification of read evidence support for all agreed-upon insertion loci, and (c) read support for the subset of agreed-upon insertion loci with Sanger sequence validations.

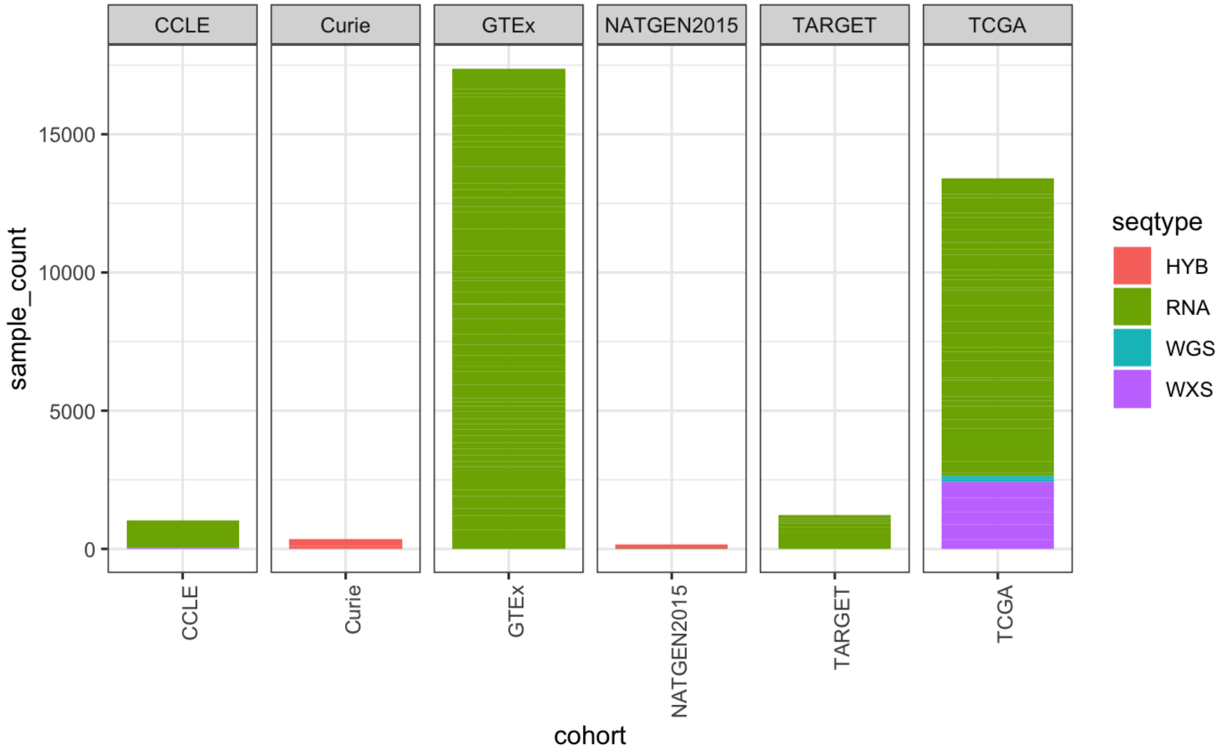


**Supplemental Figure S3: Counts of samples explored by ctat-VIF according to cohort and sequencing type.** Counts of samples are indicated for each cohort and colored according to sequencing assay: hybridization capture (HYB), RNA-seq (RNA), whole genome sequencing (WGS) or whole exome sequencing (WXS).

### Supplemental Figure S4 a-d

#### a. RNA-seq

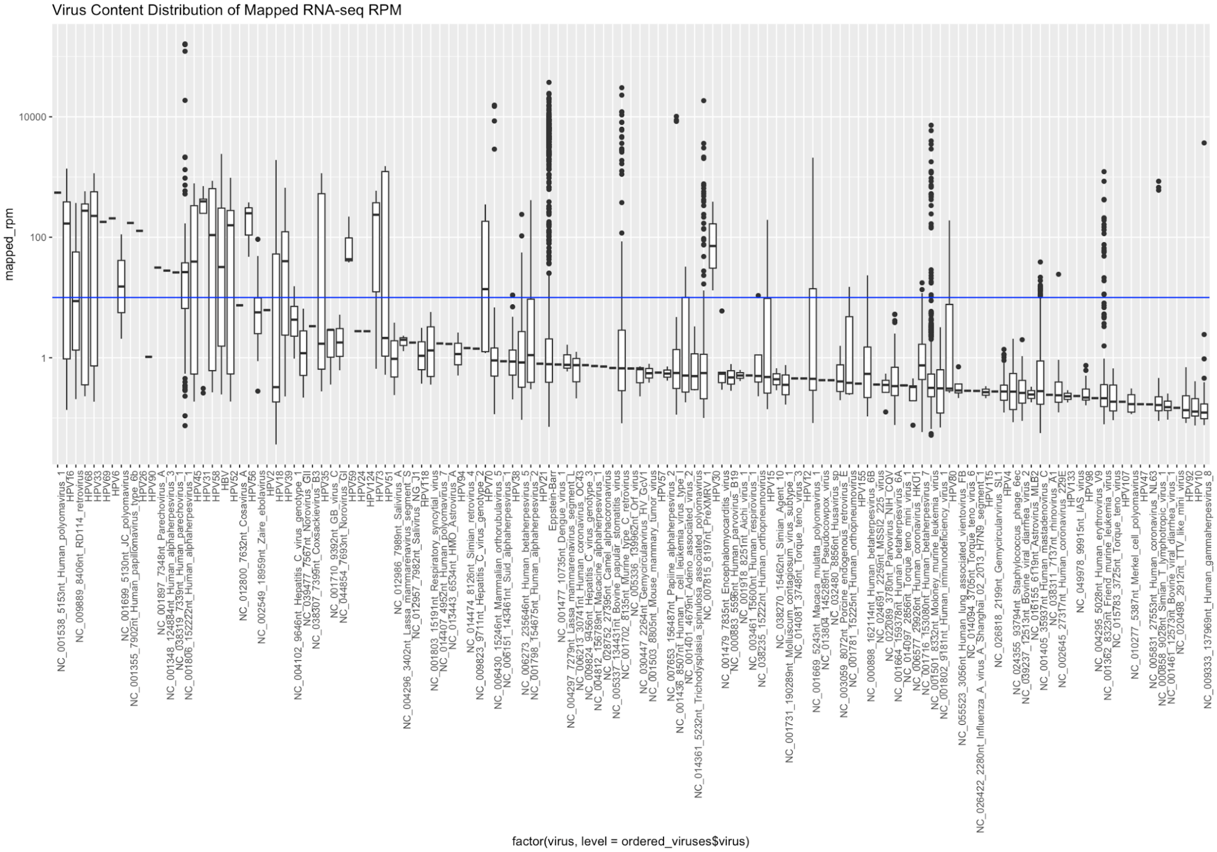

b. WXS

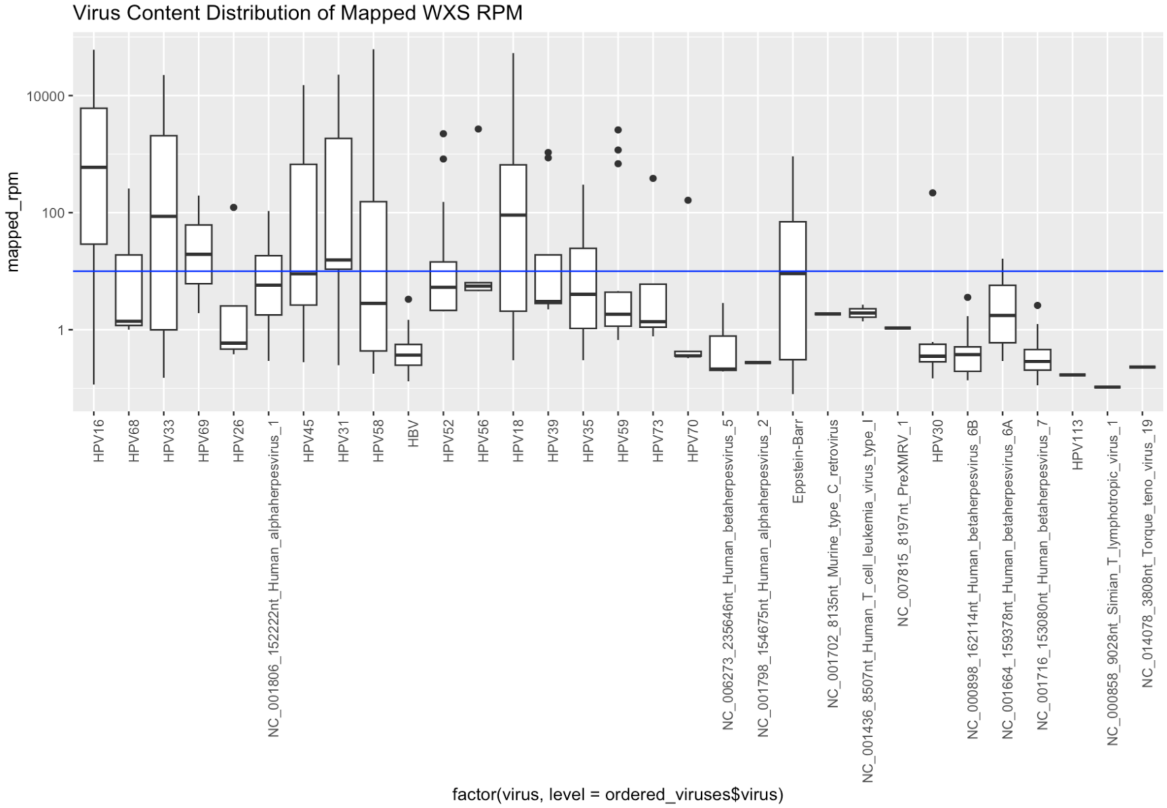
Virus Content Distribution of Mapped WXS RPM
mapped_rpm
10000
100
1
HPV16
HPV68
HPV33
HPV69
HPV26
NC_001806_152222nt_Human_alphaherpesvirus_1
HPV45
HPV31
HPV58
HBV
HPV52
HPV56
HPV18
HPV39
HPV35
HPV59
HPV73
HPV70
NC_006273_235646nt_Human_betaherpesvirus_5
NC_001798_154675nt_Human_alphaherpesvirus_2
Eppstein-Barr
NC_001702_8135nt_Murine_type_C_retrovirus
NC_001436_8507nt_Human_T_cell_leukemia_virus_type_I
NC_007815_8197nt_PreXMRV_1
HPV30
NC_000898_162114nt_Human_betaherpesvirus_6B
NC_001664_159378nt_Human_betaherpesvirus_6A
NC_001716_153080nt_Human_betaherpesvirus_7
HPV113
NC_000858_9028nt_Simian_T_lymphotropic_virus_1
NC_014078_3808nt_Torque_teno_virus_19
factor(virus, level = ordered_viruses$virus)

c. HYB-capture

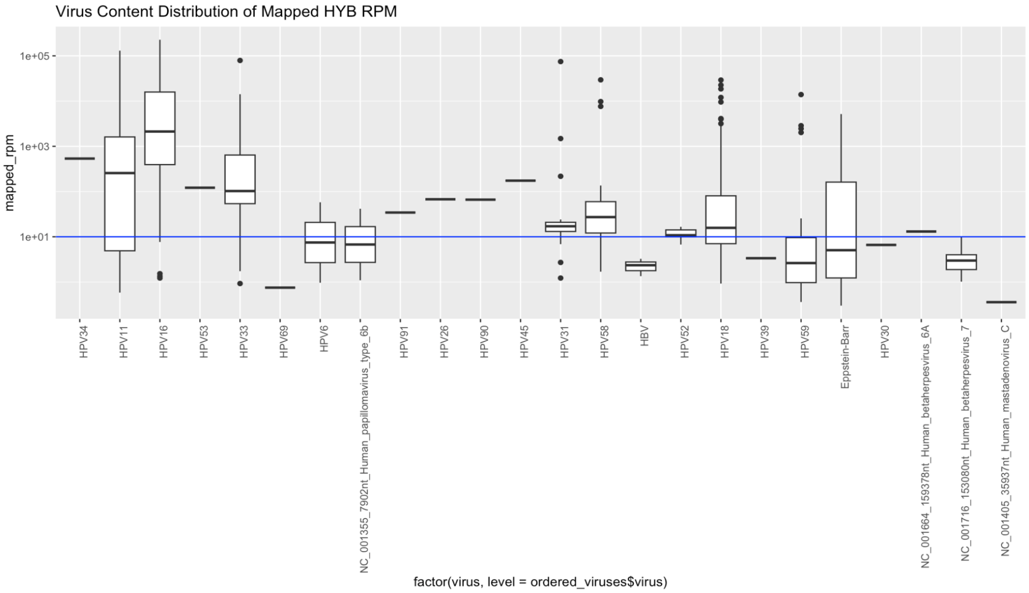


d. WGS

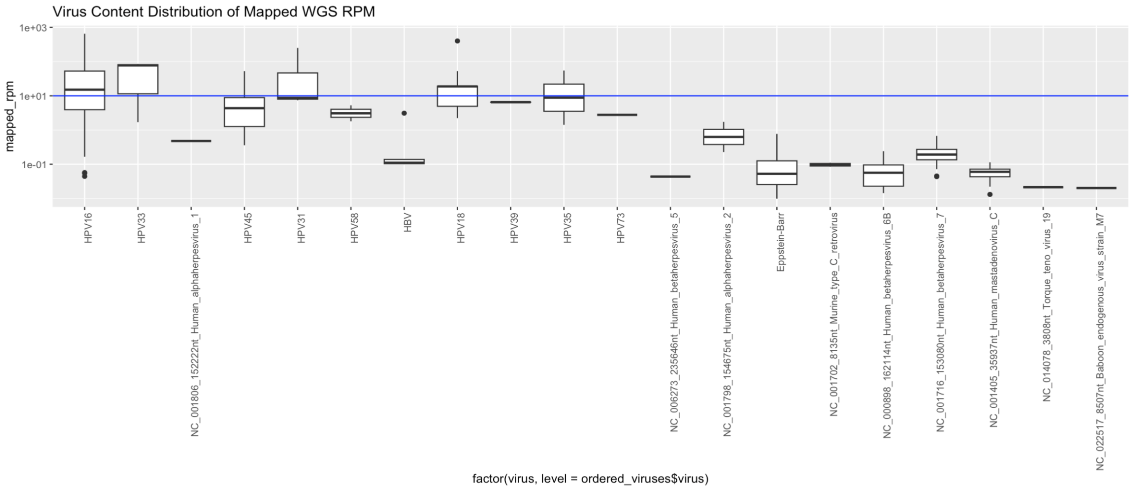


**Supplemental Figure S4: Distribution of virus mapped rpm according to virus type and sequencing type: (a) RNA-seq, (b) WXS, (c) hybrid capture, or (d) WGS.** A horizontal black line is drawn at 10 rpm, the threshold chosen to discriminate lower trace-level virus detection.

### Supplemental Figure S5

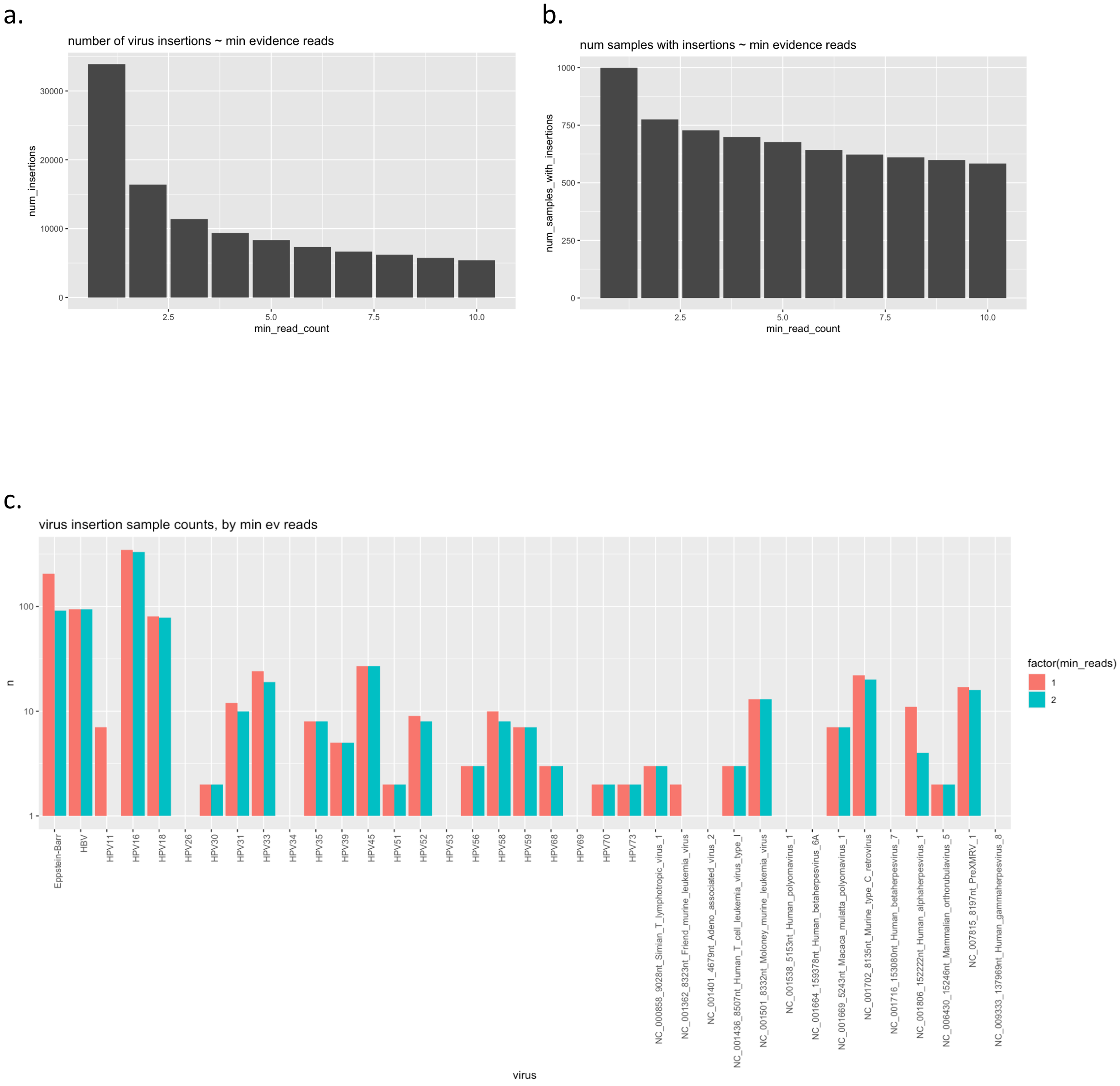

d.

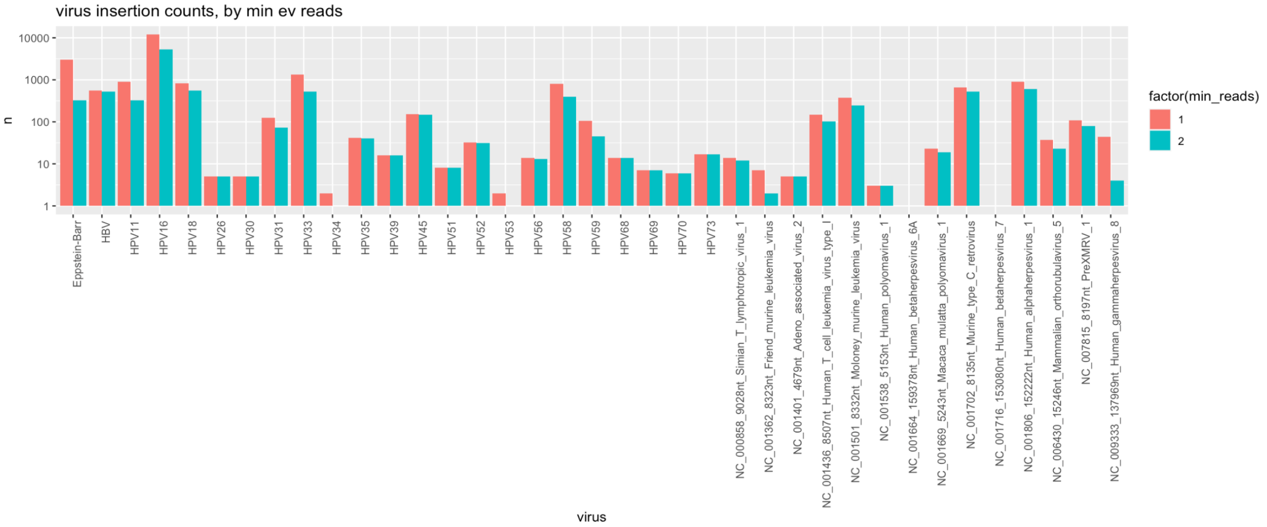


**Supplemental Figure S5: Insertion and sample counting according to minimum evidence thresholds.** (a) Number of insertions or (b) samples according to minimum insertion read evidence count threshold. (c) Counts of samples with virus insertions or (d) counts of virus insertions according to virus type and either a minimum of 1 or 2 evidence reads.

**Supplemental Figure S6 a-d**

a.

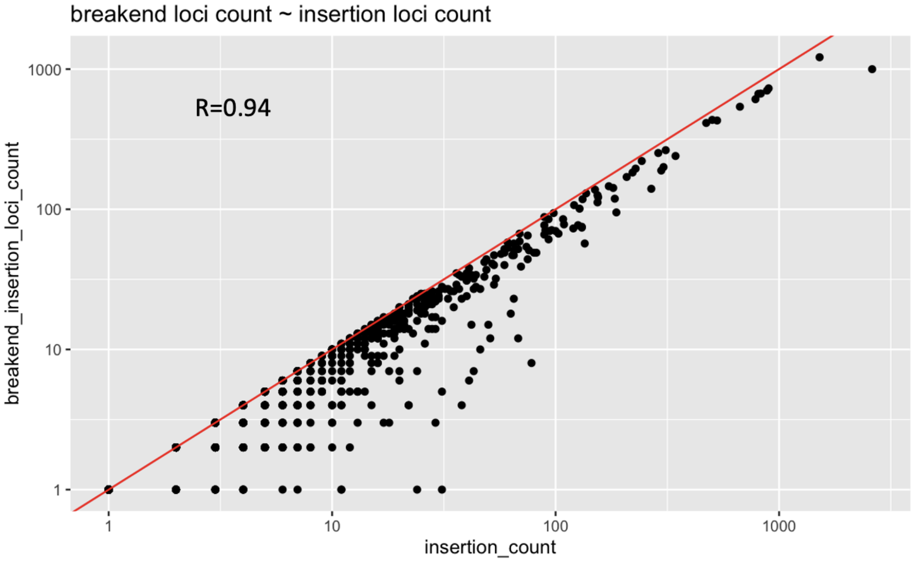

b.

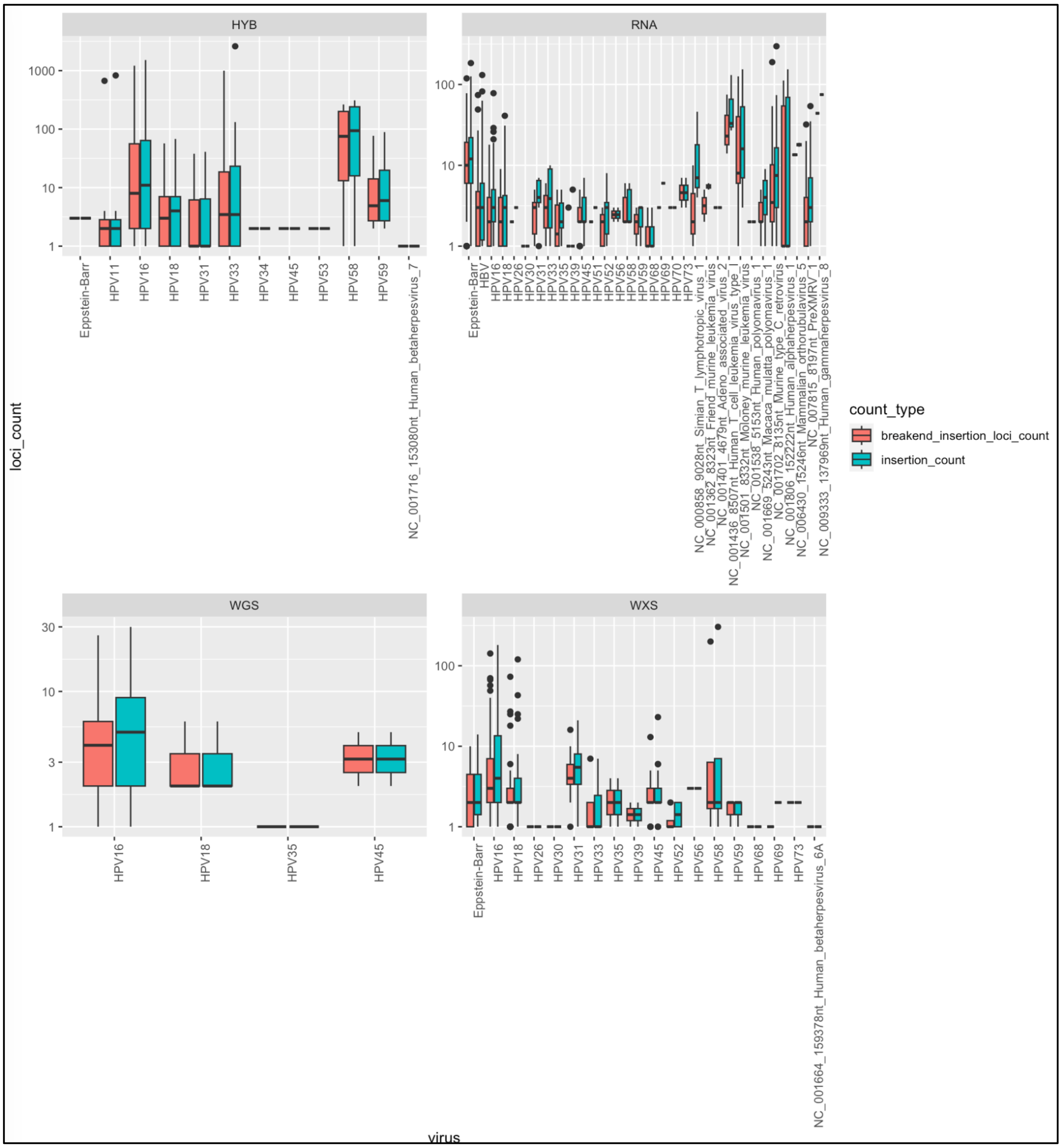

HYB
RNA
WGS
WXS
loci_count
virus
count_type
breakend_insertion_loci_count
insertion_count

c.

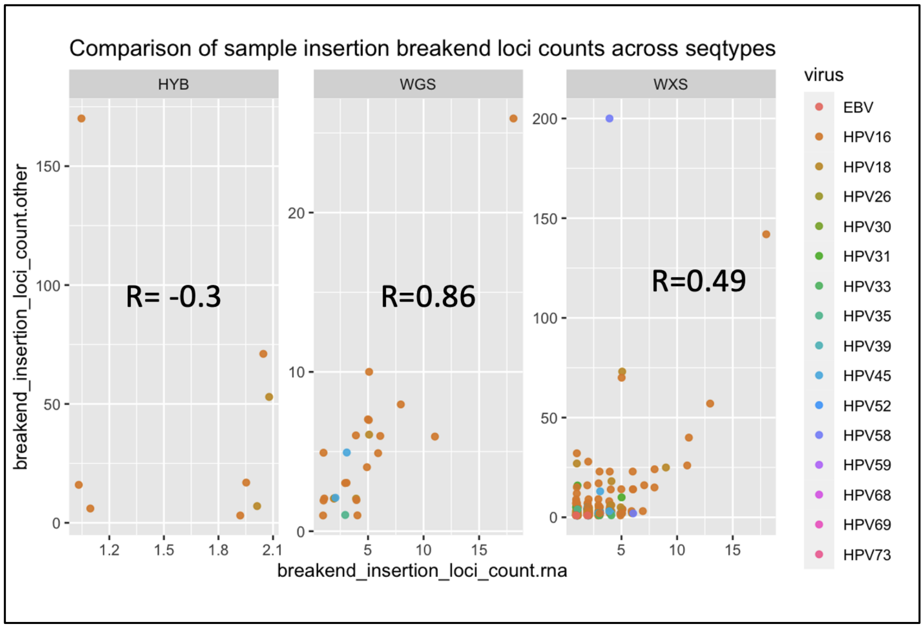


d.

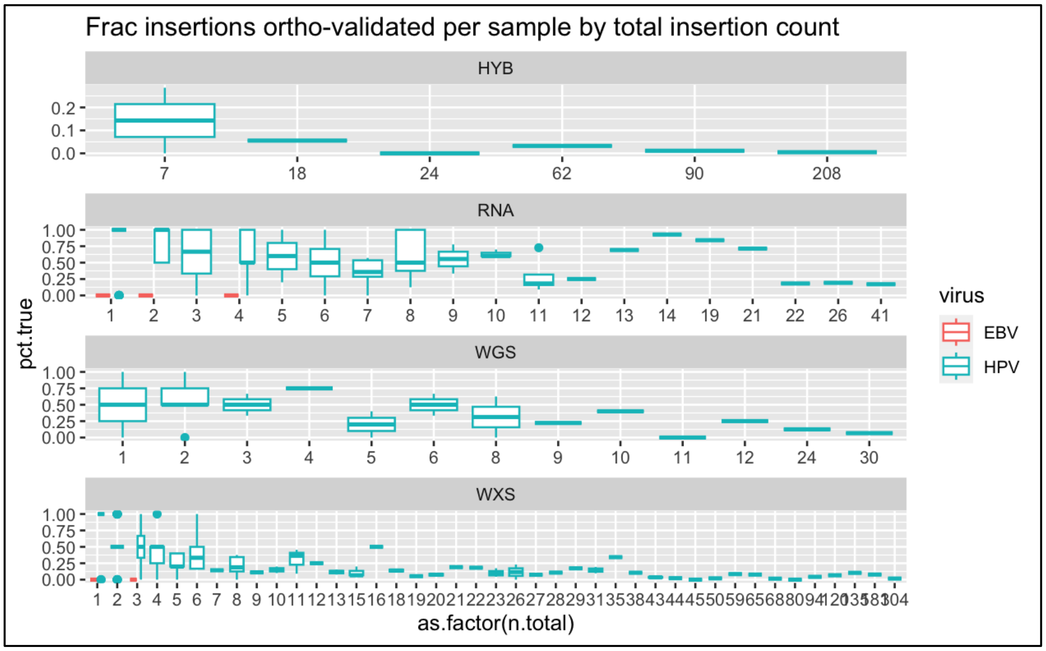


**Supplemental Figure S6: Comparison of virus insertion counts to virus break-end counts according to virus and sequencing type.** (a) Insertion counts and corresponding break-end counts are highly correlated. (b) Distribution of insertion and break-end counts according to virus (x-axis) and sequencing type (facet). (c) Number of virus insertion break-ends compared between RNA-seq and the alternative sequencing types. (d) Distribution of fractions of insertions per sample that are considered validated by RNA-seq or orthogonal sequencing type (facet) structured according to number of RNA-seq reads supporting the insertion sites.

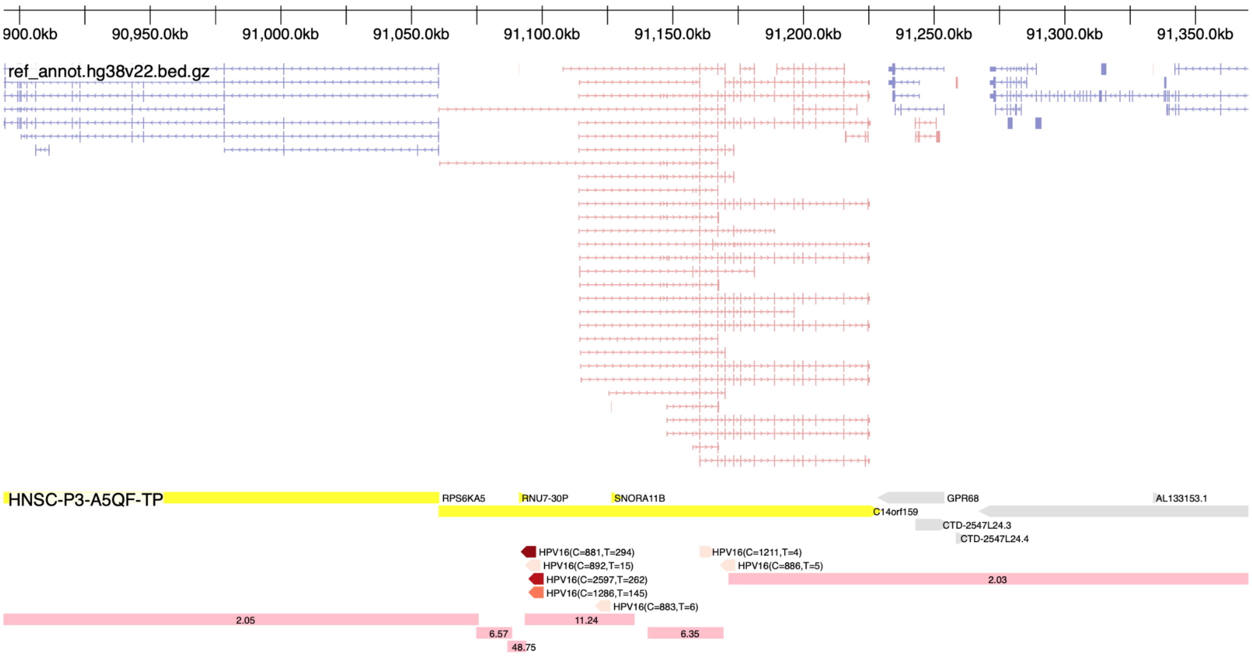


**Supplemental Figure S7: HPV16 insertions at the C14orf159 locus in head and neck tumor sample HNSC-P3-A5QF-TP coincides with a focal amplification of 49 copies**. From top to bottom: reference gene structures shown (pink=top strand, blue=bottom strand), gene spans with expression outliers indicated (yellow: top expression quantile, >= 0.95), virus insertion sites (C: viral genome breakpoint coordinate, T: total number of chimeric reads as supporting evidence), and regions with copy number indicated (pink bars).

**Supplemental Figure S8 a-c**

**a.**

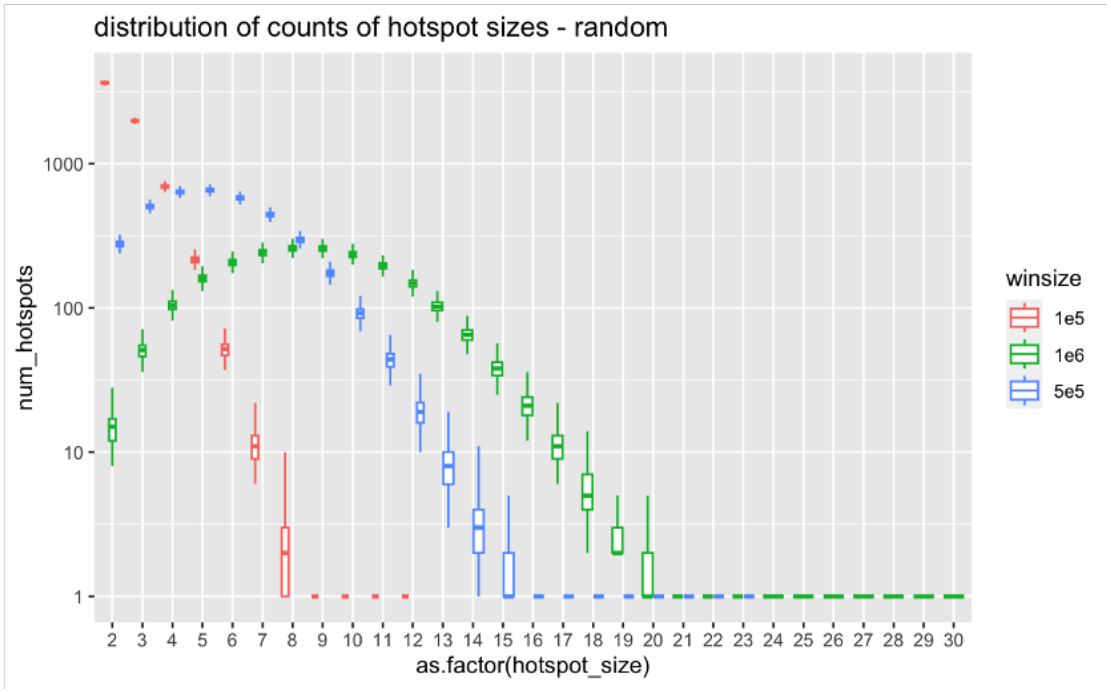

b.

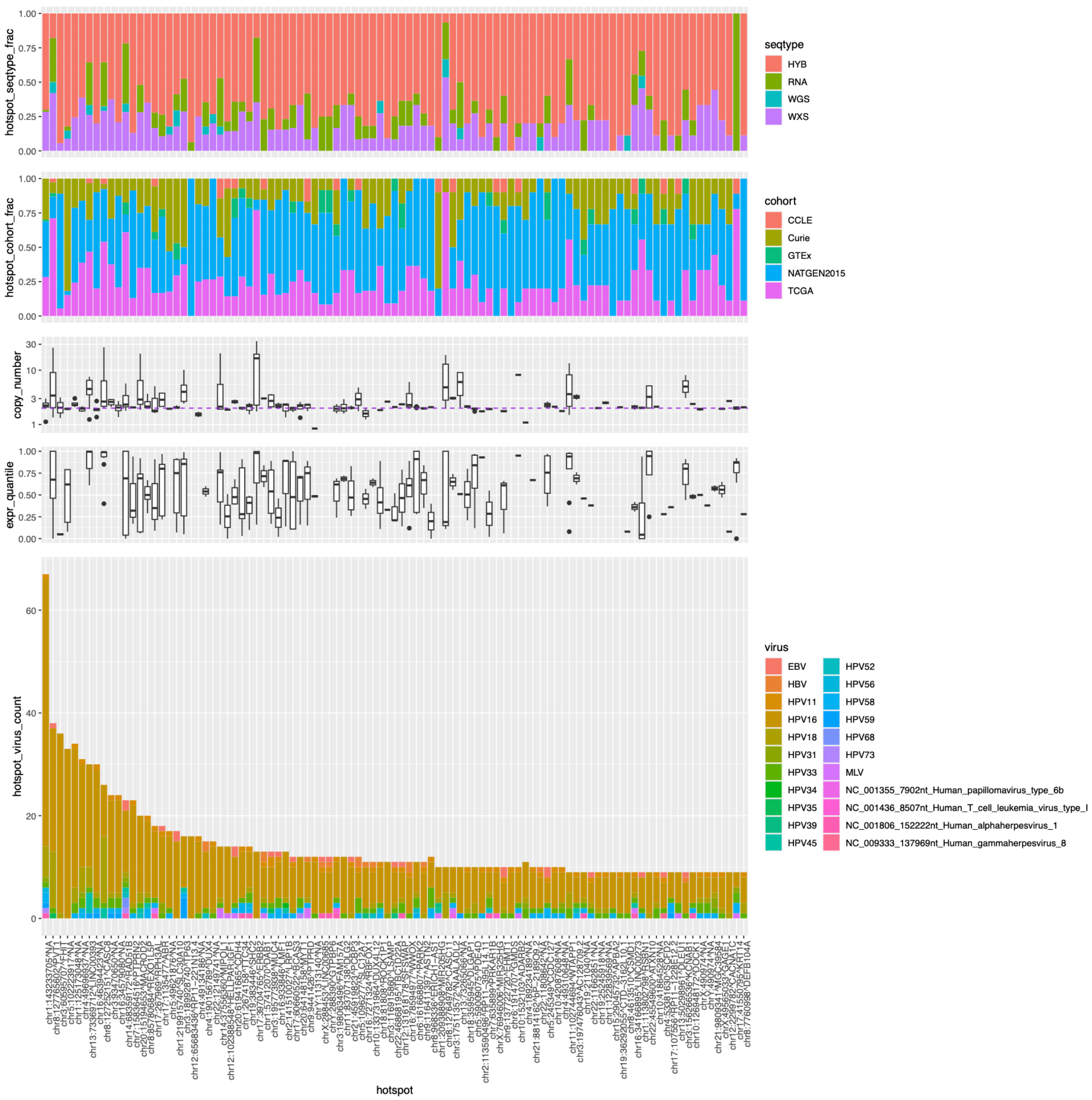
seqtype
HYB
RNA
WGS
WXS
hotspot_seqtype_frac
cohort
CCLE
Curie
GTEx
NATGEN2015
TCGA
hotspot_cohort_frac
copy_number
expr_quantile
virus
EBV
HBV
HPV11
HPV16
HPV18
HPV31
HPV33
HPV34
HPV35
HPV39
HPV45
HPV52
HPV56
HPV58
HPV59
HPV68
HPV73
MLV
NC_001355_7902nt_Human_papillomavirus_type_6b
NC_001436_8507nt_Human_T_cell_leukemia_virus_type_I
NC_001806_152222nt_Human_alphaherpesvirus_1
NC_009333_137969nt_Human_gammaherpesvirus_8
hotspot_virus_count
hotspot

c.

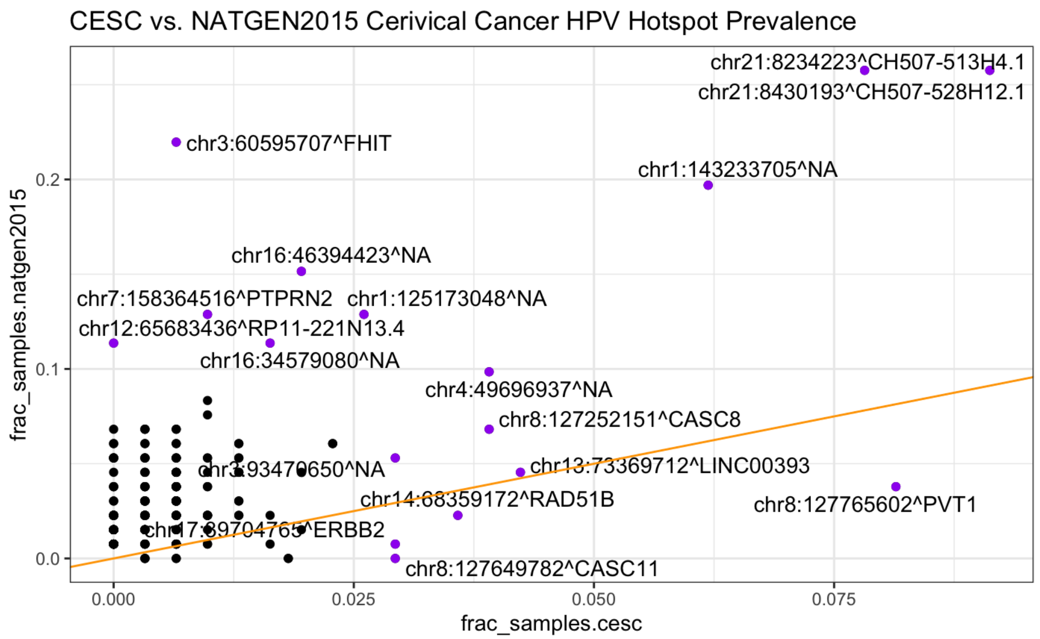


**Supplemental Figure S8: Evaluation of Viral Insertion Hotspots.** (a) Distributions of hotspot sample representation according to window size definition with random insertion sites with 10k trials. (b) HPV-enriched insertion hotspots and attributes according to top ranking hotspots. Attributes examined include (from top-to-bottom): distribution of hotspot insertion support according to sequencing modality, cohort representation, copy number at insertion sites, expression quantile for genes at insertion site, and counts of virus types at insertion sites. (c) Comparison of fraction of cervical cancers with insertions at corresponding hotspots according to Hu et al., 2015 (NATGEN2015) or from our analysis of TCGA using CTAT-VIF.

**Supplemental Figure S9 a,b**

a.

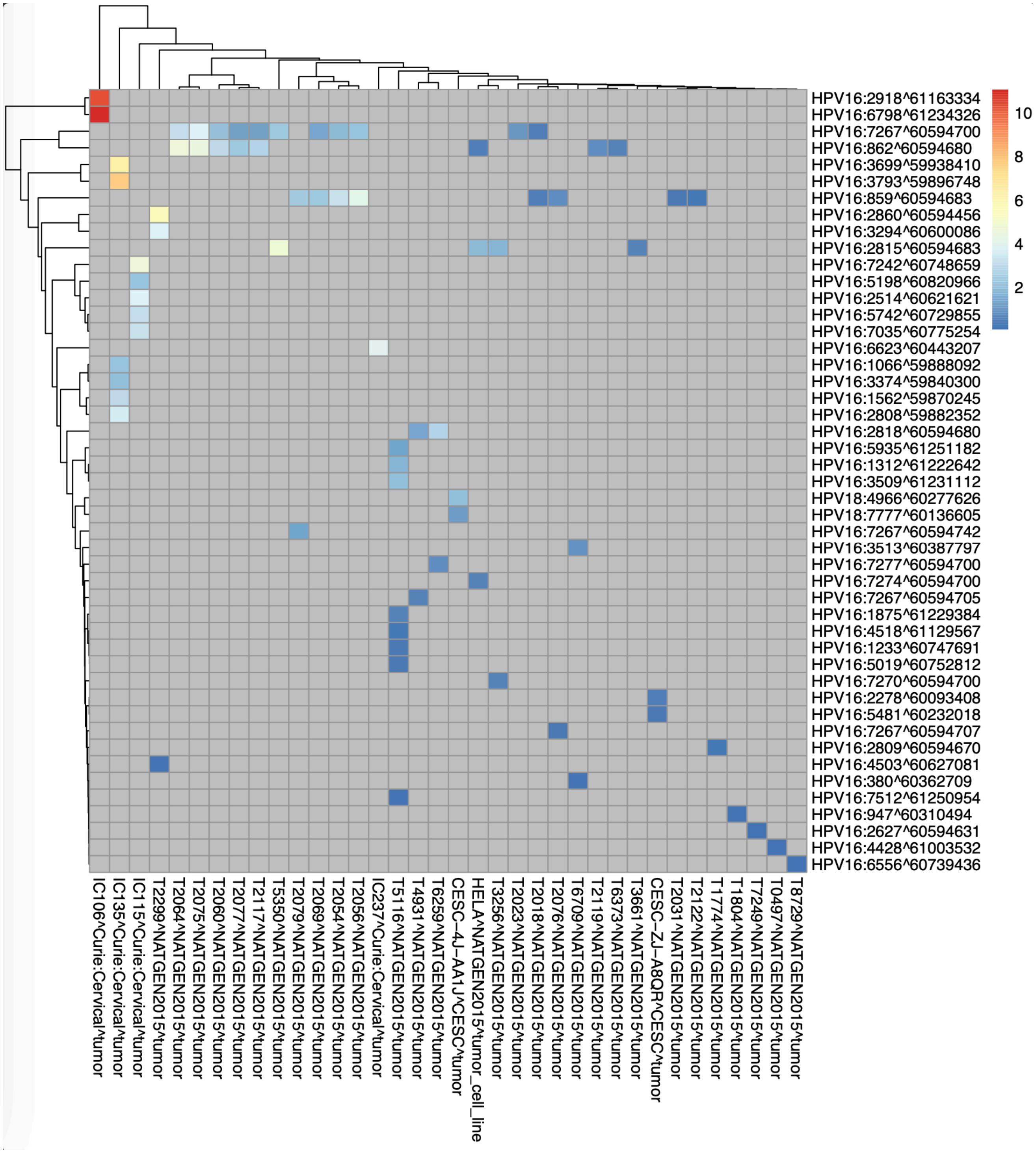

b.

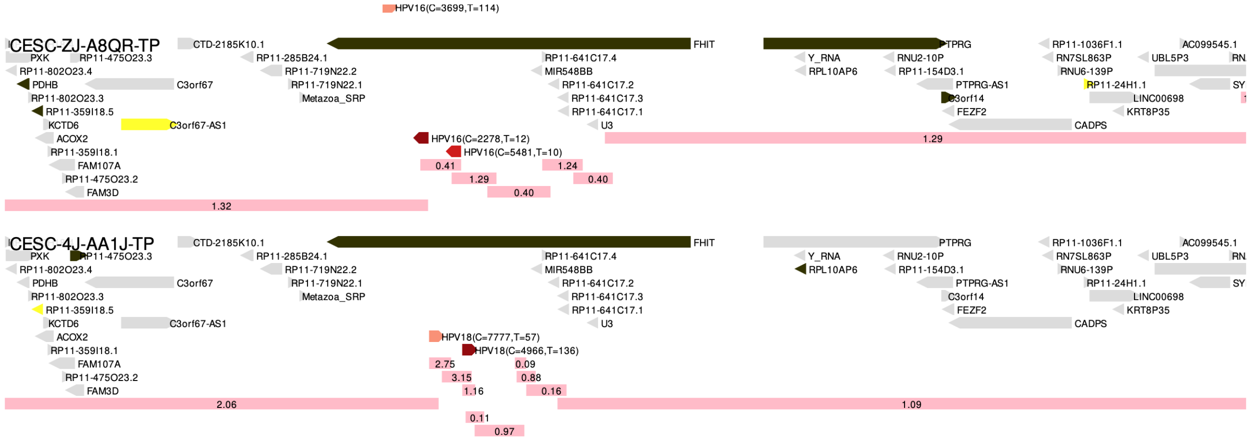


**Supplemental Figure S9: Insertions detected at the FHIT hotspot.** Several Hu et al., 2015 (NATGEN2015) cohort samples have identical insertions. Also, the HELA cell line is known to harbor HPV18 but not HPV16, yet HPV16 is detected, furthering evidence for sequencing contamination. (b) Functional impacts of insertions in two CESC samples at the FHIT locus. From top-to-bottom: genes and insertions are shown as directional spikes colored black (low expression quantile), yellow (high expression quantile) or gray (central 90% expression quantile). Virus insertion positions are shown (center) with viral genome breakpoint coordinate (C) and total read support (T) indicated. At bottom are regions of the genome with copy numbers assigned according to TCGA with copy numbers indicated in the pink rectangles.
